# Aromatic amino acid biosynthesis by a *Lotus* Aldolase impacts root hair development and symbiotic associations

**DOI:** 10.1101/2022.10.13.511206

**Authors:** Jesús Montiel, Euan K. James, Ivette García-Soto, Dugald Reid, Selene Napsucialy-Mendivil, Joseph G. Dubrovsky, Luis Cárdenas, Jens Stougaard

## Abstract

- Legume roots can be symbiotically colonized by arbuscular mycorrhizal (AM) fungi and nitrogen-fixing bacteria. In *Lotus japonicus*, the latter occurs intracellularly by the cognate rhizobial partner *Mesorhizobium loti* or intercellularly with the *Agrobacterium pusense* IRBG74 strain. Although these symbiotic programs show distinctive cellular and transcriptome signatures, some molecular components are shared.
- In this study, we demonstrate that *Aldolase1*, the first enzyme in the biosynthetic pathway of aromatic amino acids (AAA), plays a critical role in root hair development and for AM and rhizobial symbioses in *Lotus*.
- Two homozygous mutants affected in *Aldolase1* (*ald1*-1 and *ald1*-2) show drastic alterations in the root hair morphology, associated with a progressive disruption of the actin cytoskeleton. The altered root hair structure was prevented by chemical and genetic complementation.
- Both *ald1*-1 and *ald1*-2 show significant reductions in rhizobial infection (intracellular and intercellular), nodule organogenesis and AM colonization. RNAseq analysis of *ald1*-2 roots suggested that these phenotypes are associated with downregulation of several cell wall related genes, and with an attenuated symbiotic signalling. This work provides robust evidence that links AAA metabolism to root hair development and successful symbiotic associations.

## Introduction

Legume roots establish interactions with beneficial microorganisms in the rhizosphere through complex chemical dialogues. Arbuscular mycorrhizal symbiosis (AMS) and nitrogen-fixing symbiosis (RNS) are two well-known examples, which provide phosphorus and nitrogen sources, respectively, to the host (Oldroyd, 2013). In the model legume *Lotus japonicus* (*Lotus*), two modalities of root colonization exist in the RNS, depending on the bacterial partner (Montiel *et al*., 2021; Quilbe *et al*., 2022). The cognate rhizobial partner *Mesorhizobium loti* synthesizes and releases lipochito-oligosaccharide nodulation factors (NFs) signalling molecules, after perception of flavonoid compounds secreted by roots (Bek *et al*., 2010; Shimamura *et al*., 2022). Recognition of these compounds activates the symbiotic signalling pathway that allows rhizobial infection and nodule development. *M. loti* attaches to the root hair tip, inducing its curling and trapping the bacteria within infection pockets, where a tubular structure called an infection thread (IT) is formed. In this process, the rhizobial partner colonizes the root cell layers intracellularly, through root hair ITs, which advances towards the cortex, where a nodule primordium, generated by reactivation of the cell division in the root cortex, is formed. The bacteria are released from the ITs into the nodule cells as organelle-like structures, called symbiosomes, differentiating into nitrogen-fixing bacteroids (Roy *et al*., 2020). However, our group recently showed that *Lotus* can be also infected intercellularly by the *Agrobacterium pusense* IRBG74 strain (Cummings *et al*., 2009; Mitra *et al*., 2016). In this entry mode, IRBG74 induces massive root hair curling and twisting, but no root hair ITs are formed; instead, the bacteria pass between the epidermal root cells, forming cortical infection pockets. The migration proceeds both intra- and intercellularly into the nodule cells, releasing the bacterial content from transcellular ITs or intercellular peg-infection structures (Montiel *et al*., 2021).

Under phosphate deficiency, strigolactones secreted by the roots are perceived by AM fungi, promoting spore germination and hyphal branching. These responses favour the physical interaction of the fungal hyphae with the root epidermal cells, giving rise to the formation of the hyphopodium (Bonfante & Genre, 2010). The fungal hyphae penetrate the epidermal cells, reaching the cortex, where they differentiate into branched hyphae called arbuscules. In these structures, carbon and phosphorous sources are exchanged between the symbionts (MacLean *et al*., 2017). Molecular genetic studies have identified important players involved in the signalling pathway that allow the mutualistic associations. In *Lotus*, the NFs produced by *M. loti* (Bek *et al*., 2010) are perceived at the root hair plasma membrane by the nod factor receptors *NFR1* and *NFR5* (Madsen *et al*., 2003; Radutoiu *et al*., 2003; Murakami *et al*., 2018; Bozsoki *et al*., 2020; Gysel *et al*., 2021). This recognition is followed by the rhizobial infection, which is coordinated by different transcription factors such as nodule inception (Nin; Schauser et al., 1999; Marsh et al., 2007), nodule signalling pathway Nsp1/Nsp2 (Kalo *et al*., 2005; Heckmann *et al*., 2006), Cyclops (Yano *et al*., 2008; Singh *et al*., 2014), and ERF required for nodulation (Ern1; (Cerri *et al*., 2012; Kawaharada *et al*., 2017)). Similarly, in the last decades, a growing list of molecular components necessary for IT growth and development have been identified (Roy *et al*., 2020). The intercellular colonization of IRBG74 in *Lotus* roots shows evident cellular and transcriptome differences, with distinct genetic requirements with respect to the root hair IT formation process. However, there is a core of symbiotic genes that are indispensable for any modality of rhizobial infection (Montiel *et al*., 2021; Quilbe *et al*., 2022), which belong to the common symbiotic signalling pathway (CSSP) and also play essential roles in the AMS (Stracke *et al*., 2002; Levy *et al*., 2004; Yano *et al*., 2008).

The root hairs are extensions of specialized epidermal cells with polarized growth and a tubular shape, which increase the surface area for nutrient acquisition. In *Arabidopsis thaliana*, the characterization of a large collection of mutants affected during root hair development and emergence indicates that these processes are regulated by a sophisticated network that encompass various molecular functions (Shibata & Sugimoto, 2019). Root hairs also play a crucial role during the RNS, in the early signalling pathway and in the intracellular colonization of rhizobia (Downie, 2014). Actin cytoskeleton rearrangements and cell wall modifications in the root hairs are necessary to facilitate the formation and progression of ITs (Cardenas *et al*., 2003; Yokota *et al*., 2009; Hossain *et al*., 2012; Zepeda *et al*., 2014; Qiu *et al*., 2015). Several regulators of root hair development also participate in the establishment of symbiotic associations with rhizobia and AM fungi (Karas *et al*., 2005; Lei *et al*., 2015; Ke *et al*., 2016; Liu *et al*., 2020; Karas *et al*., 2021).

The functioning of the nitrogen-fixing nodule and arbuscules is regulated by a large battery of metabolic genes. In these symbiotic organs, several enzymes of diverse metabolic pathways modulate the flux of carbon, phosphorous and nitrogen sources between the root cells and the microsymbionts (Udvardi & Poole, 2013). However, transcriptome analyses of legume roots inoculated with compatible rhizobia and AM fungi show that reprogramming of metabolic processes also occurs at early stages of the symbiotic associations (Manthey *et al*., 2004; Deguchi *et al*., 2007; Handa *et al*., 2015; Fonseca-Garcia *et al*., 2021). Despite the evident activation of several metabolic routes during the colonization and organogenesis programs in the mutualistic interactions of legumes, their role has been poorly explored. In this study, we found that expression of a *Lotus Aldolase* (*LjAldolase1*) gene was associated with several stages of the RNS and root hair development. *Aldolase1* encodes a Phospho-2-dehydro-3-deoxyheptonate aldolase (also named DAHP synthase), the first enzyme of the shikimate pathway, which leads to the biosynthesis of the aromatic amino acids (AAA) phenylalanine, tyrosine and tryptophan (Lemaitre *et al*., 2014). These amino acids serve as precursors of various metabolites, including phytohormones and cell wall components (Tzin & Galili, 2010; Geng *et al*., 2020; Simpson *et al*., 2021). The characterization of two homozygous mutant alleles disrupted in the *LjAldolase1* revealed a pivotal role of this gene in root hair development and the establishment of symbiotic associations with nitrogen-fixing bacteria and AM fungi.

## Material and Methods

### Germination, nodulation kinetics and genotyping

The genotypes used in this study belong to the genetic background of *L. japonicus* accession Gifu (Handberg & Stougaard, 1992). For germination, the seedcoat was mechanically removed with sandpaper, and the subsequent surface was sterilized with sodium hypochlorite (3%) for 10-15 minutes and washed 3-5 times with sterile distilled water to remove traces of chlorine. The imbibed seeds were transferred to square Petri dishes with damp paper and incubated at 21°C for germination. For nodulation assays, three-to-five-day-old seedlings were transferred to square Petri dishes with a 1.4% agar slant supplemented with ¼ B&D solution (Broughton & Dilworth, 1971), which provides minimal requirements for plant growth without nitrogen source that favour the RNS. The agar was covered with autoclaved filter paper and inoculated with *M. loti* R7A or IRBG74 (1 mL of bacterial culture per plate: OD600 = 0.05). The experiment was conducted in a growth room with photoperiod (16/8 h) and was temperature controlled at 21°C. Using a stereomicroscope, the number of white and pink nodules were recorded weekly at 1-6 weeks post inoculation (wpi). The LORE1 lines 30100225 and 30141487, affected in the *LjAldolase1* gene, were obtained from the LORE1 mutant collection (Urbanski *et al*., 2012; Malolepszy *et al*., 2016) and genotyped to obtain homozygous mutants with allele-specific primers, following the database guidelines (Mun *et al*., 2016).

### Root phenotyping and pharmacological complementation

Gifu, *ald1*-1 and *ald1*-2 seeds were surface sterilized as mentioned above and transferred to square Petri dishes with an 0.8% agar slant supplemented with 0.2x MS medium and 1% sucrose, which allows optimal growth and contains a nitrogenous source. For the chemical complementation, 50 and 100 µM of L-Phenylalanine, L-Tryptophan and L-Tyrosine were added to the agar individually or in a mixture (50 and 100 µM each AAA). The experiment was conducted in a growth chamber with controlled temperature and photoperiod. The root growth dynamics were analysed by measuring the root length daily in the different genotypes, from the radicle emergence up to 11 dpg. The lengths of the apical meristem and root hairs were measured from images obtained with a stereomicroscope on 10 dpg plants.

The actin cytoskeleton of 4 dpg seedlings of different genotypes was visualized with epifluorescence microscopy on root segments fixed with Alexa-Phalloidin (Yokota *et al*., 2009). The dynamics of filamentous actin plus ends was monitored in live root hairs of Gifu, *ald1*-1 and *ald1*-2 seedlings, mounted carefully in adapted Petri dishes with 1 ml of Fahraeus medium containing 4 µl of Cyt-Fl probe (Molecular Probes; 2.5 µM) (Zepeda *et al*., 2014). Cells incubated with Cyt-Fl were excited at 484 nm, and emission was collected at 530 nm (20 nm band pass). All filters used were from Chroma Technology, and image acquisition and analysis were carried out using MetaMorph/MetaFluor software (Universal Imaging, Molecular Devices).

### *In silico* analyses and RNAseq

The nucleotide and peptide sequences of LjAldolase1, LjAldolase2 and LjAldolase3 were extracted from the *Lotus* genome browser (Mun *et al*., 2016; https://lotus.au.dk/genome/). The presence of the signal peptide in the amino acid sequences was predicted with TargetP – 2.0 (https://services.healthtech.dtu.dk/service.php?TargetP-2.0). Aldolases from other plant species were obtained through protein BLAST in Phytozome (Goodstein *et al*., 2012), using as a query the three *Lotus* Aldolases. The amino acid sequences were aligned and bootstrapped (NJ tree, 1,000 iterations) by ClustalX 2, and the tree was visualized with Dendroscope (Huson & Scornavacca, 2012). The accession numbers and annotations of Aldolase sequences are indicated in Table S1

The *ald1*-2 seeds were germinated, and the seedlings were inoculated on plates with IRBG74 or mock-treated (with water), following the protocol mentioned above. The root segments susceptible to intercellular infection (elongation and maturation zone) were cut and frozen in liquid nitrogen at 5 dpi for total RNA isolation. The RNA integrity and concentration were determined by gel electrophoresis and Nanodrop, respectively. DNA contamination was removed through DNAse treatment. Library preparations using randomly fragmented mRNA were performed by IMGM laboratories (Martinsried, Germany) and sequenced in paired-end 150-bp mode on an Illumina NovaSeq 6000 instrument. A decoy-aware index was constructed for Gifu transcripts using default Salmon parameters and reads were quantified using the validate Mappings flag (Salmon version 0.14.1; (Patro *et al*., 2017)). Normalized expression levels and differential expression testing were calculated with the R-package DESeq2 version 1.20 (Love *et al*., 2014) after summarizing gene level abundance using the R-package tximport (version 1.8.0).

### Root infection phenotyping and nodule histology

The different genotypes were inoculated with the *M. loti*-LacZ strain, and the roots were harvested at 1 wpi for histochemical staining with X-Gal and recording the IT progression within the root tissues. The intercellular infection was analysed as previously described (Montiel *et al*., 2021). Gifu and *ald1*-2 roots were collected at 3 wpi with IRBG74, incubated for 1 min in a solution for surface disinfection (0.3% of chlorine and 70% EtOH), and then washed five times with distilled water. The total DNA was extracted from individual roots, adjusted to 10 ng/µl and used as template for qPCR to evaluate the IRBG74 *NodA* abundance as previously reported (Montiel *et al*., 2021) using the delta Ct method (Pfaffl, 2001).

Three-week-old nodules were detached from Gifu and *ald1*-2 plants inoculated with *M. loti* or IRBG74 and preserved in a fixative solution (0.1 M sodium cacodylate pH 7, 2.5% glutaraldehyde). Fixed nodule slices were embedded in acrylic resin and sectioned for light microscopy and for TEM. For light microscopy analysis, the nodule semi-thin sections (1 µm thickness) were stained with toluidine blue, while for TEM the ultrathin sections (80 nm thickness) were stained with uranyl acetate (James & Sprent, 1999; Madsen *et al*., 2010).

### Constructs for promoter activity and subcellular localization

The predicted promoter (2 kb upstream from the start codon) and coding sequence (1,617 bp) of *LjAldolase1* were obtained from the *L. japonicus* Gifu genome (Kamal *et al*., 2020) in the *Lotus* database (Mun *et al*., 2016), and synthetized with 5′overghangs for goldengate cloning with compatible vectors. Promoter, CDS, fluorescent reporters and terminator modules were assembled and cloned into a pIV10 vector (Stougaard, 1995), suitable for goldengate technology. *Agrobacterium rhizogenes* strain AR1193 was transformed with the pAldolase1::tYFP-nls and pAldolase1::Aldolase1-YFP constructs used to induce transgenic hairy roots in Gifu and *ald1*-2 at 6 dpg, following a standardized protocol (Hansen *et al*., 1989). Three weeks after infection, the main root was removed, and the plants with hairy roots were transferred to square Petri dishes with ¼ B&D agar or plastic magenta boxes containing LECA substrate. Depending on the construct, the transgenic roots were mock-treated or inoculated with either *M. loti*-DsRed (Kelly *et al*., 2013) or IRBG74-DsRed (Montiel *et al*., 2021) for inspection by confocal microscopy.

### Confocal microscopy of fluorescent reporters and symbiotic colonization

The confocal microscopy analysis was carried out in a Zeiss LSM780 microscope with excitation laser/emission filter (nm) settings adjusted to the fluorescent markers: autofluorescence, 405/408– 498 nm; YFP, 514/517–560 nm; GFP, 514/517-540; DsRed, 561/517–635 nm. Gifu and *ald1*-2 plants were inoculated onto plates with the fluorescent-labelled *M. loti*-DsRed (Kelly *et al*., 2013) and IRBG74-DsRed strains. For AM analysis, the tissue was incubated for 4 h in EtOH (70%) at room temperature, then transferred to KOH (20%) for 2-3 days and washed three times with water. Later, the root segments were treated with HCl (0.1 M) for 1-2 h, and the solution was replaced by PBS containing 1 µg/ml^-1^ of WGA-Alexa Fluor 488 (Torabi *et al*., 2021). The root segments inoculated with the fluorescent bacteria, the transgenic roots harbouring the pAldolase::tYFP-nls and pAldolase::Aldolase1-YFP constructs, and the tissue stained with WGA-Alexa Fluor 488 were mounted onto microscope slides for observation.

The analysis of root hairs after pharmacological treatments was performed on root segments from plants germinated and grown in media supplemented with AAA, as mentioned above. The tissue was cleared using the ClearSee-adapted protocol (Kurihara *et al*., 2015). For cell wall visualization, the last step of the protocol consisted of 40 min incubation in ClearSee solution supplemented with 0.1% of Calcofluor White, for which Fluorescent Brightener 28 disodium salt (Sigma-Aldrich F3397) was used. The roots were analysed under a confocal laser scanning microscopy setup built around a Zeiss Axiovert 200M microscope (Oberkochen, Germany) that consisted of a high-speed galvo-resonant scanner for visible wavelengths (SCANVIS), a 405-nm laser OBIS 405nm LX 100mW laser system, 495nm longpass dichroic, 440/40, nm bandpass emission filter, photomultiplier tube (PMT) modules (standard sensitivity), a Z-Axis piezo stage with controllers, and ThorImageLS 4.0 software. All parts from Thorlabs, Inc (Newton, NJ, USA).

### Arbuscular mycorrhization quantification

Four-day-old seedlings of Gifu and *ald1*-2 were placed between two discs of cellulose membrane filters (0.22 µm pore size) with 50-100 spores of *R. intraradices* (Symbiom®), previously resuspended in Long-Ashton solution (Hewitt & Smith, 1975) and the carrier substrate provided by the manufacture. The filters with the inoculated plants were covered with autoclaved sand within Magenta boxes. At 4 and 6 wpi, the root system was detached from the plants and cleared as follows: 2% KOH (1 h at 90 °C), 3-5 washes with distilled water, 2% HCl (30-60 min), staining with trypan blue solution (1:1:1, lactic acid, glycerol and water; 15-60 min at 90°C) and washed with 50% glycerol. Stained root segments were mounted onto microscope slides, and different fungal structures were recorded with the help of an optical microscope, following the protocol described by Trouvelot et al. (1986).

## Results

### An *Aldolase* is induced during the *Lotus*-IRBG74 symbiosis

We have recently shown that IRBG74 induces a distinctive transcriptome response during the intercellular infection of *Lotus* roots (Montiel *et al*., 2021). Among the regulated genes, LotjaGi1g1v0143000 encoding a Phospho-2-dehydro-3-deoxyheptonate aldolase (hereafter referred as *Aldolase1*) was the most abundant transcript (Fig. 1a). Aldolase1 is the first enzyme of the shikimate pathway (Fig. S1a) involved in the biosynthesis of AAA (Tzin & Galili, 2010), which suggests an unexpected role for amino acid homeostasis in the RNS. Supporting this notion, *Aldolase1* is highly expressed in NF-treated root hairs, nodule primordia and mycorrhized roots (Fig. S1c). A further investigation of the gene in the *Lotus* database (https://lotus.au.dk/) shows that *Aldolase1* contains five exons and four introns (Fig. 1b). To determine whether *Aldolase1* belongs to a gene family, a BLAST was performed in the *Lotus* genome (Kamal *et al*., 2020). Two homolog genes were identified, LotjaGi1g1v0239000 and LotjaGi6g1v0351200, which were named *LjAldolase2* and *LjAldolase3*, respectively (Fig. 1b). Unlike *LjAldolase1, LjAldolase2* and *LjAldolase3* were not significantly induced by IRBG74 at any timepoint studied (Fig. S1b) and showed lower expression levels in root hairs, nodule primordia, and mycorrhized roots (Fig. S1c). The deduced amino acid sequence of *Aldolase1* encodes a peptide of 539 amino acids, with a predicted plastid signal peptide (Fig. S2). Similarly, Aldolase2 and Aldolase3 contain an N-terminal plastid-targeting sequence. The protein sequence of these three *Lotus* enzymes is highly conserved (Fig. S2), sharing >80% of their identities. To investigate the phylogeny of *Lotus* Aldolases, a phylogenetic tree was constructed using full-length peptide sequences. In this analysis, the *Lotus* Aldolases were compared with homolog sequences found in the legumes *Aeschynomene evenia, Cicer arietinum, Glycyrrhiza uralensis, Medicago truncatula* and *Phaseolus vulgaris*. In addition, their phylogenetic distribution was evaluated with their counterparts in several non-legumes (monocots and dicots) with fully sequenced genomes, i.e. *Oryza sativa, Sorghum bicolor, Arabidopsis thaliana, Solanum lycopersicum* and *Solanum tuberosum*. The Aldolase family was composed of four members in the monocots *O. sativa* and *S. bicolor*, while in dicots it ranged between two and four members (Fig. 1c). Particularly, legumes of the inverted repeat-lacking clade (IRLC) contained only two Aldolases, except for *G. uralensis* that possess three (Fig. 1c). The different Aldolases clustered in subclades that reflect the phylogeny of the species.

**Fig. 1.**
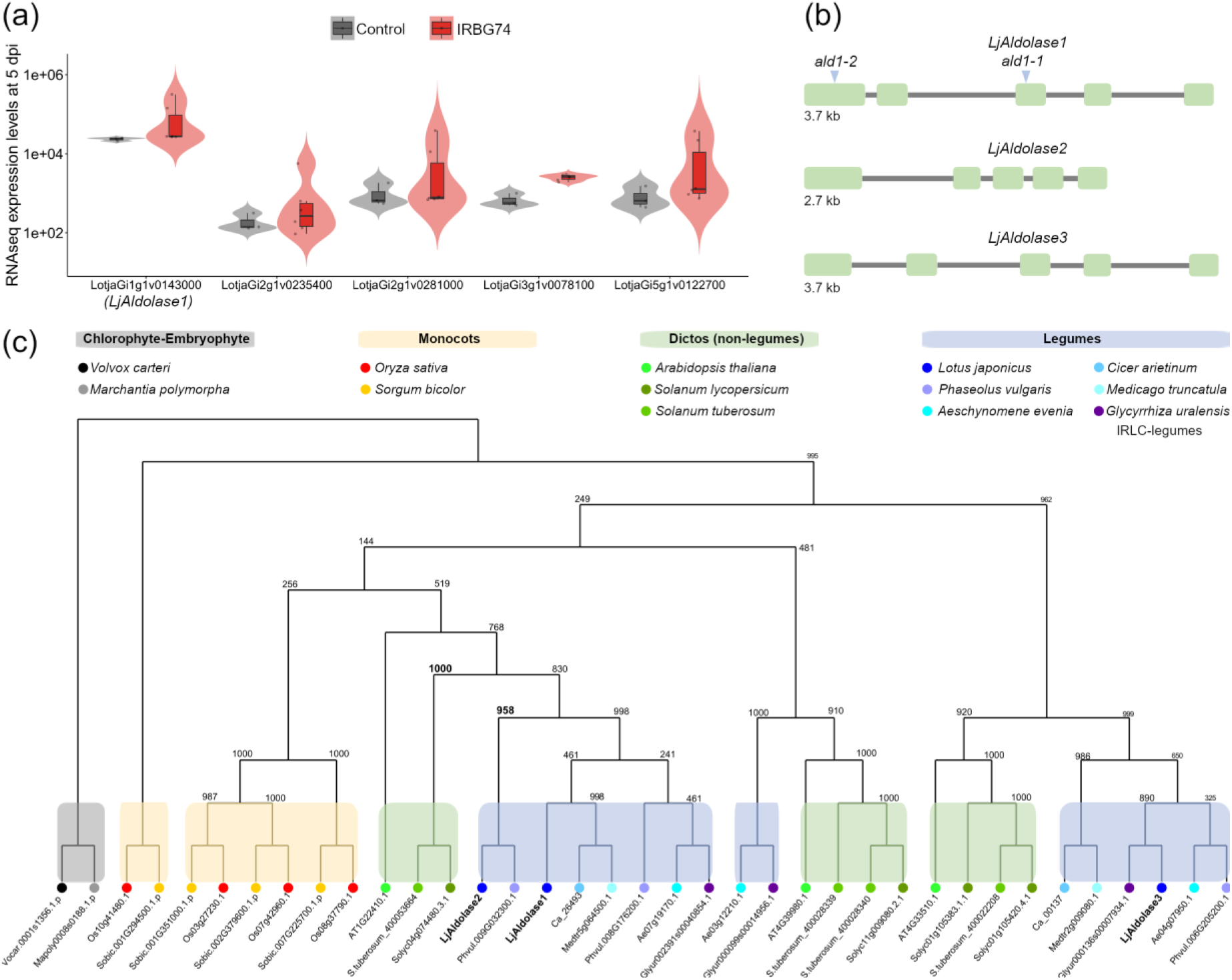
Highly expressed genes during the *Lotus*-IRBG74 symbiosis, gene structure and phylogeny of Aldolases. (a) violin dot plots show the expression levels of upregulated genes with the highest transcript abundance in Gifu roots at 5 dpi with IRBG74. Data calculated from RNAseq information (Montiel et al. 2021). (b) graphic representation of the exon (green) and intron (grey line) composition in the *LjAldolase1* (LotjaGi1gv0143000), *LjAldolase2* (LotjaGi1g1v0239000) and *LjAldolase3* (LotjaGi6g1v0351200) genes. Blue arrowheads indicate the retrotransposon insertions in the *ald1*-1 (30141487) and *ald1*-2 (30100225) alleles. (c) phylogenetic relationship of Aldolases in various legumes and non-legumes (monocots and dicots). The chlorophyte (*Volvox carteri*) and embryophyte (*Marchantia polymorpha*) Aldolases were included as root. Bootstrap values are shown at the nodes.

**Fig. S1.**
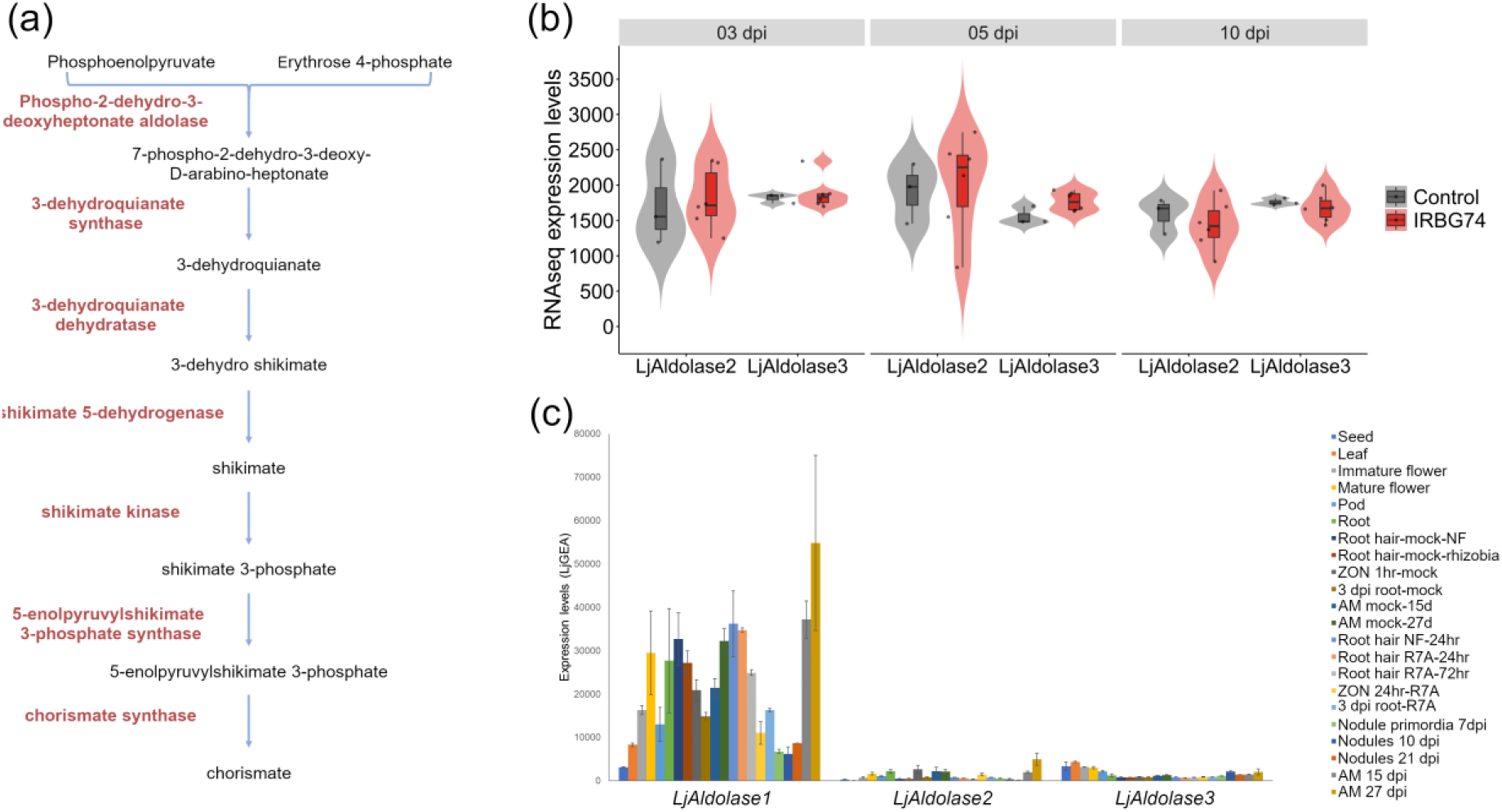
Expression profile of *Lotus* Aldolases in different tissues and conditions. (a) compounds (black) and enzymes (red) involved in the shikimate pathway. (b) expression levels of *LjAldolase2* and *LjAldolase3* extracted from RNAseq data of *Lotus* roots at 3, 5 and 10 dpi with IRBG74 or mock-treated (control). (c) expression levels of *LjAldolase1, LjAldolase2* and *LjAldolase3* in different tissues, organs and in response to different stimuli. Data extracted from the *Lotus japonicus* Expression Atlas (https://lotus.au.dk/expat/).

**Fig. S2.**
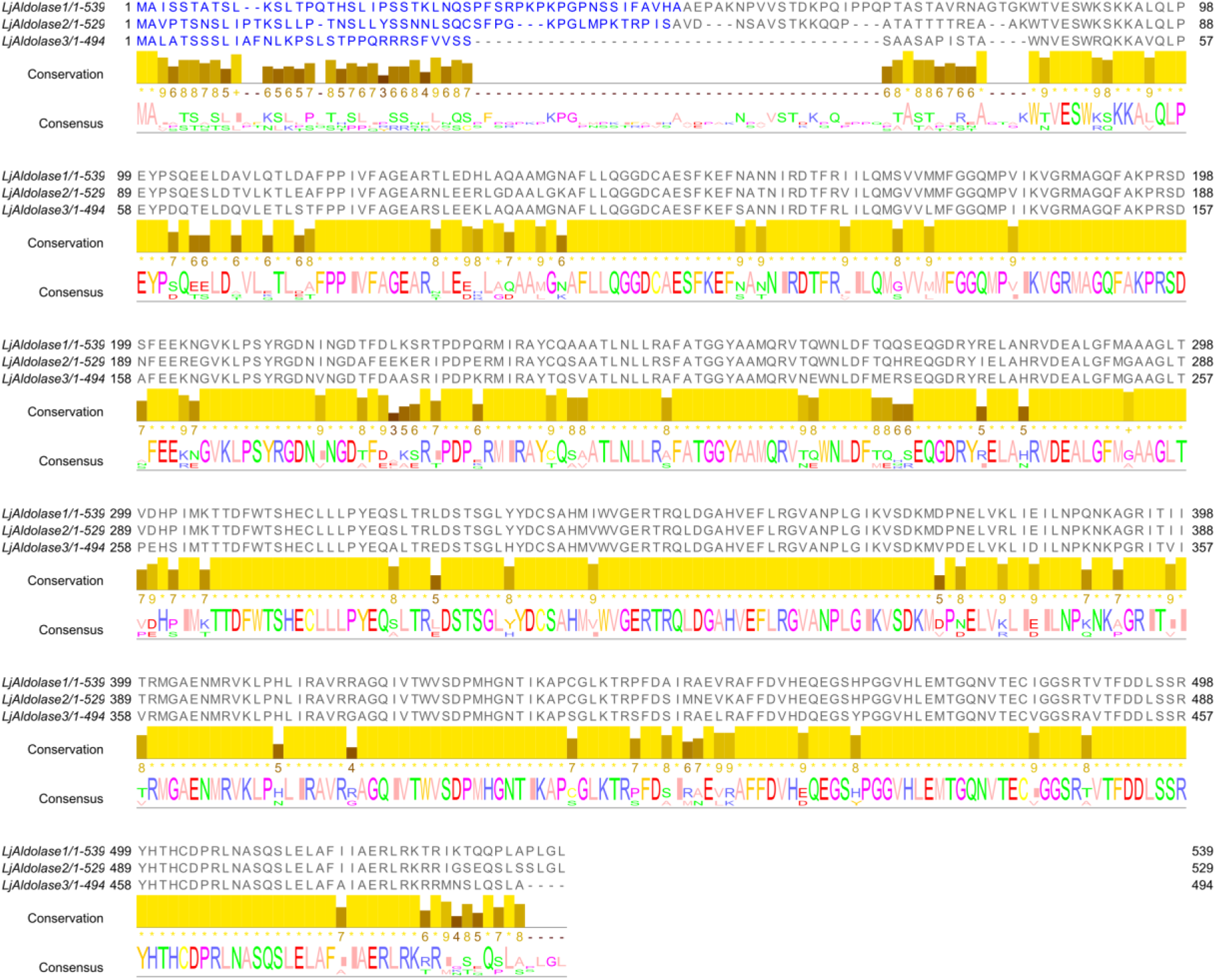
Multiple sequence alignment of *Lotus* Aldolases. Multiple alignment of deduced full peptide sequences of LjAldolase1, LjAldolase2 and LjAldolase3. In blue, residues of the predicted plastid signal peptide.

### High promoter activity of *Aldolase1* in root tissues and during RNS

Transcriptome profile sources indicates that *Aldolase1* plays a role in the RNS of *Lotus*. To explore the cellular expression pattern, we monitored the activity of the *Aldolase1* 2 kb promoter sequence fused to the triple yellow fluorescent protein with a nuclear localization signal (*pAldolase1::tYFP-nls*). Strong expression of the fluorescent reporter was detected in the apical region of the root, growing root hairs and emerging lateral root primordia of uninoculated roots (Fig. 2a-c). Likewise, after *M. loti*-DsRed and IRBG74-DsRed inoculation, *pAldolase1::tYFP-nls* was detected at high levels in the epidermal and cortical cells adjacent to intracellular and intercellular infection sites, respectively (Fig. 2d,e). An elevated promoter activity remained during nodule organogenesis (Fig. 2f), which indicates the participation of *Aldolase1* during nodule development.

**Fig. 2.**
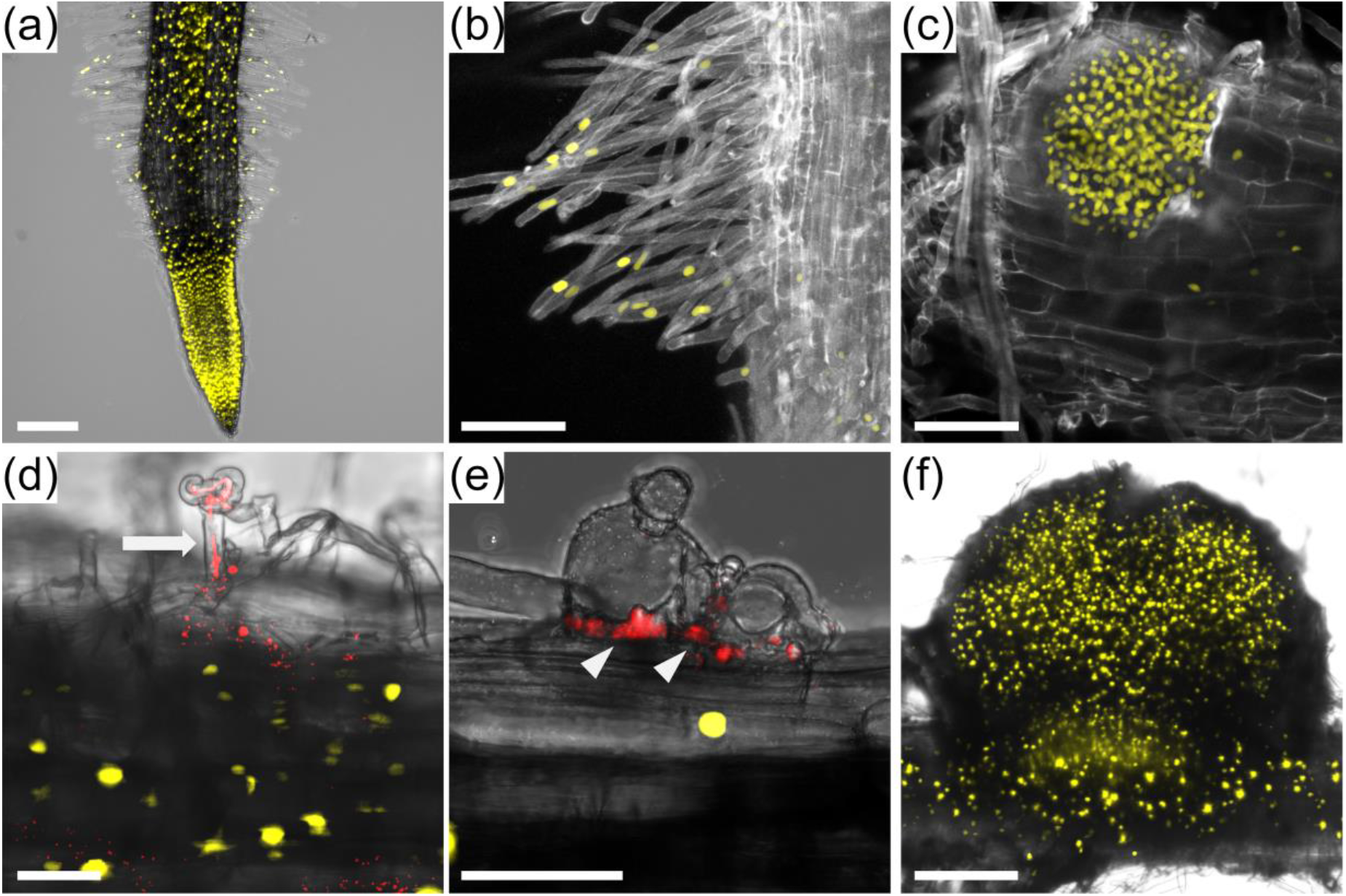
Promoter activity of *LjAldolase1* by confocal microscopy in *Lotus* roots. Expression of *Aldolase1* promoter was evaluated in uninoculated (a-c) and inoculated (d-f) *Lotus* transgenic roots harbouring the *pAldolase1::tYFP-nls* construct. (a), root tip. (b), root hairs. (c), lateral root primordium. Strong promoter activity in cells surrounding the root hair IT (d), the intercellular infection (e) and nodule organogenesis at 1 wpi with *M. loti*-DsRed (d) and 2 wpi with IRBG74-DsRed (e and f). Arrow and arrowheads indicate the IT and intercellular infection, respectively. Scale, 50 µm (d and e), 100 µm (b and c), and 200 µm (a and f).

### *LjAldolase1* is crucial for root hair growth

To further investigate the potential role of *Aldolase1* in the symbiotic associations of *Lotus*, two homozygous mutant alleles, *ald1*-1 and *ald1*-2, were obtained from the LORE1 mutant collection (Urbanski *et al*., 2012; Malolepszy *et al*., 2016) and contained retrotransposon insertions in the third and first exon, respectively (Fig. 1b). The evidence obtained on the expression profile of *Aldolase1* led us to first explore the root phenotype in the *ald1*-1 and *ald1*-2 mutants. Both the primary root length and the root apical meristem (RAM) length in 10 dpg plants was not significantly different in the two mutant alleles, when compared to wild type Gifu (w. t.) plants (Fig. S3a-c). Similarly, the root growth dynamics in the *ald1*-1 and *ald1*-2 mutants was comparable to that recorded for Gifu roots at 1 to 11 dpg (Fig. S3d). However, the root hair development was dramatically altered in the *aldolase1* mutant alleles (Fig. 3a). Although the root hairs seem to emerge normally in the differentiation zone of the *ald1*-1 and *ald1*-2 mutants, they undergo a progressive swelling, forming balloon-like structures that eventually collapse (Fig. 3a). Unlike the root hairs of Gifu plants that reached up to 1 mm length, the root hairs *ald1*-1 and *ald1*-2 mutants barely reached 400 µm, reflecting the drastic impact of their abnormal development (Fig. 3b). Considering that the mutant and wild type roots grew at similar growth rates and that changes in root hair length started to be detected at 1.5-2 mm from the root tip (Fig 3b), the data suggest that root hair growth in the mutants was arrested soon after root hair bulge formation. This phenotype is consistent with the results obtained on the subcellular localization and promoter activity of *Aldolase1*, demonstrating a direct link between regulation of AAA biosynthesis and root hair development.

**Fig. S3.**
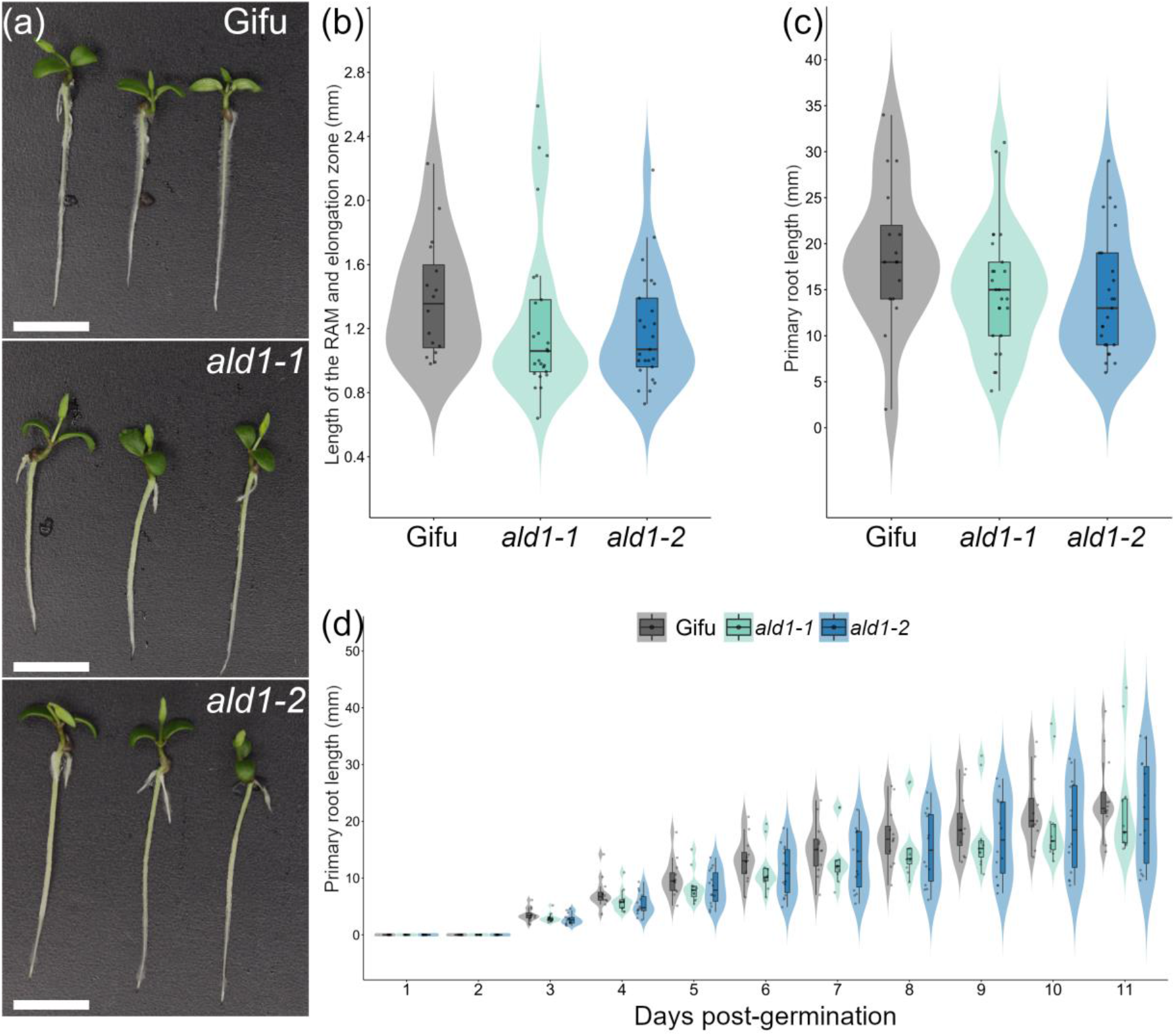
Root growth phenotype of *ald1*-1 and *ald1*-2 mutants. Representative images (a), length of the root apical meristem (RAM) with the elongation zone (b), and length of the primary root (c) in 10 dpg Gifu, *ald1*-1 and *ald1*-2 plants, grown in MS medium. (d) root growth dynamics of Gifu, *ald1*-1 and *ald1*-2 at 1-10 dpg. Violin boxplots: center line, median; box limits, upper and lower quartiles; whiskers, 1.5× interquartile range; points, individual data points. No significant differences were obtained for the different parameters when compared by Student′s t-test, between Gifu w. t. (n= 12) and the *ald1*-1 (n= 11), and *ald1*-2 (n= 14) mutants. Scale, 1 cm.

**Fig. 3.**
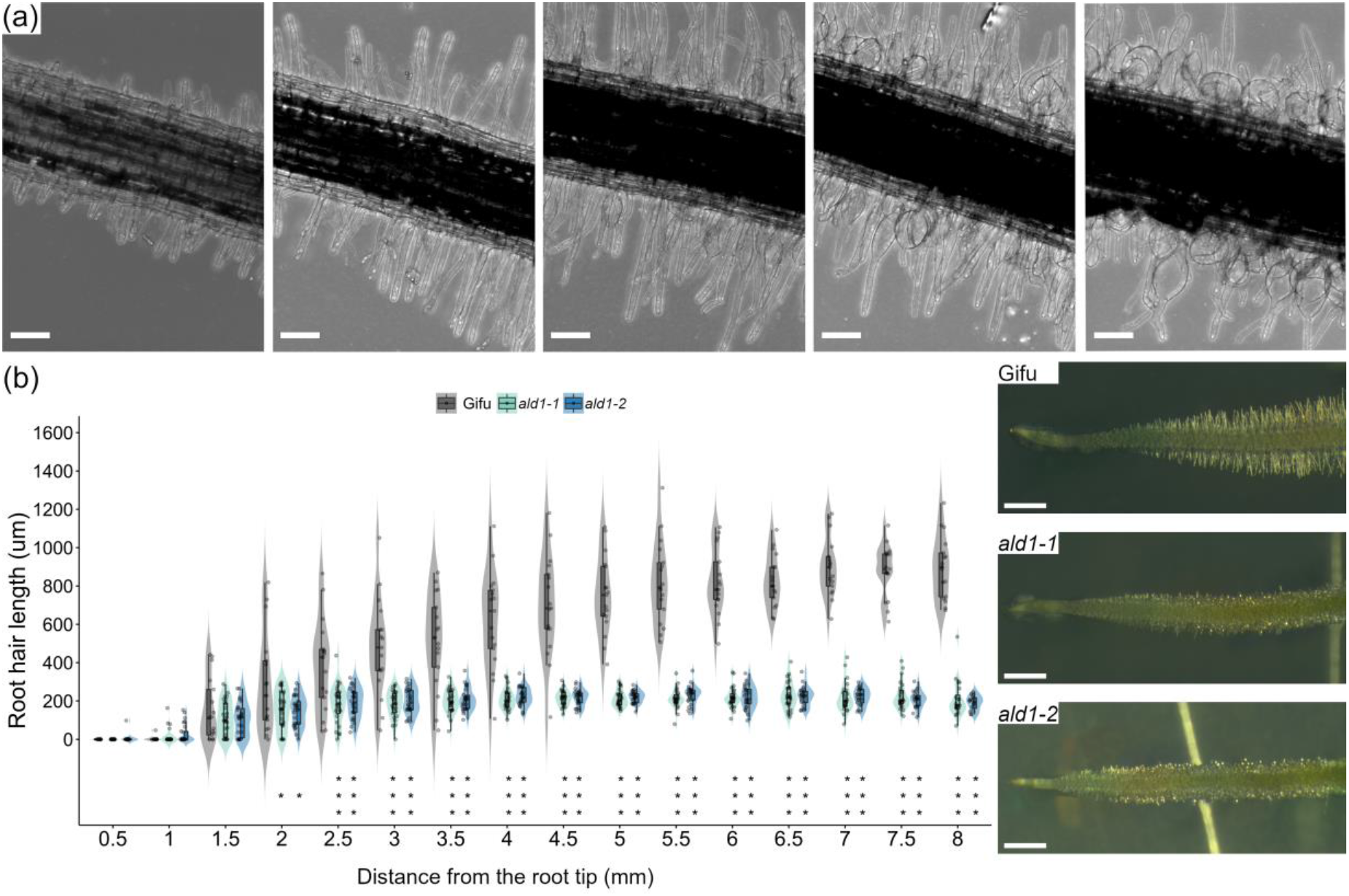
Phenotype and growth profile of root hairs in the *ald1*-1 and *ald1*-2 mutants. (a), representative images of the progressive deterioration in the root hair morphology of the *ald1*-1 mutant (the same phenotype was observed for the *ald1*-2 mutant). (b), violin dot plots of the maximal root hair length recorded at different distances from the root tip in Gifu, *ald1*-1 and *ald1*-2 roots. Panels of representative images for the different genotypes are included. Student’s t test of root hair length between Gifu (n= 17) and the *ald1*-1 (n= 27), and *ald1*-2 (n= 25) mutants. *p < 0.05; ***p < 0.001. Scale bar 100 µm (a) and 1mm (b), respectively. Violin boxplots: center line, median; box limits, upper and lower quartiles; whiskers, 1.5× interquartile range; points, individual data points.

### Actin cytoskeleton is severely disrupted in root hairs of the *aldolase1* mutants

The drastic alteration in the root hair morphology of the *ald1*-1 and *ald1*-2 mutants prompted us to study this phenomenon in more detail by analysing their actin cytoskeleton dynamics. The actin microfilaments, visualized by Alexa-Phalloidin staining, formed long bundles parallel to the long axis of w. t. root hairs (Fig. 4a). Similarly, these structures were observed in the root hairs of the *ald1*-1 and *ald1*-2 mutants with an early and mild swelling (Fig. 4b,c,f). However, the progressive alterations in the root hair morphology were accompanied by a gradual disruption of the actin cytoskeleton (Fig. 4d,e,f). To further understand the dramatic changes in the cytoskeleton organization, the actin polymerization sites were monitored in growing root hairs incubated with fluorescently-labelled cytochalasin D (Cyt-Fl). In Gifu root hairs, the major fluorescent signal was detected in the apical region, reflecting the fast-growing (plus) ends of microfilaments (Fig. 4g). In deformed root hairs of the *ald1*-1 and *ald1*-2 mutants, the fluorescence was considerably reduced in the apical region and, interestingly, sites of actin polymerization were associated to subapical regions (Fig. 4h). This displacement of actin polymerization sites seems to be linked to the misorientation of actin bundles (Fig. 4f) and would explain the root hair phenotype. Our results suggest that the progressive loss in the organization of the cytoskeleton and root hair morphology is caused by changes in the growth dynamics of actin microfilaments.

**Fig. 4.**
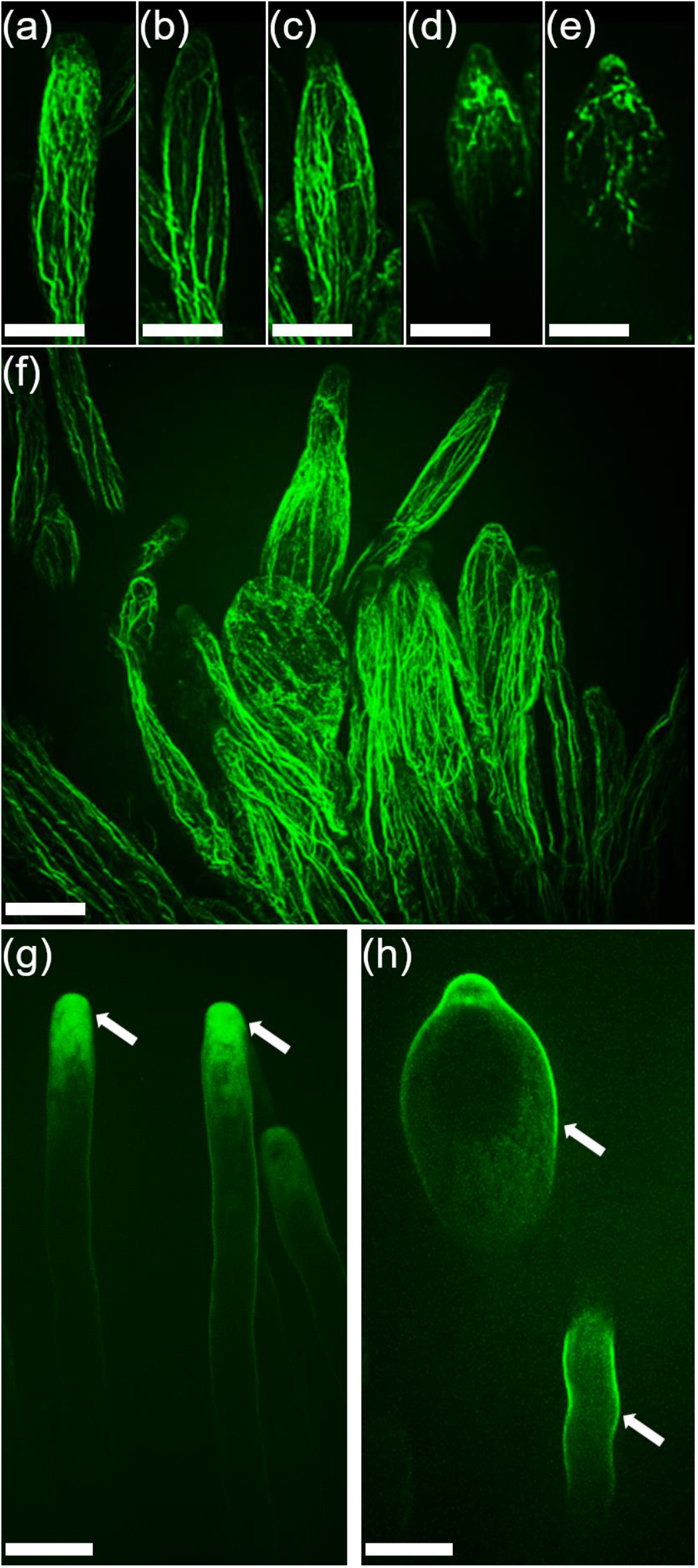
Actin cytoskeleton dynamics of *Lotus* root hairs. Epifluorescence microscopy images of Gifu (a, g), *ald1*-1 (b-e, h) and *ald1*-2 (f) root hairs stained with Alexa-Phalloidin (a-f) or fluorescently labelled cytochalasin D (g, h) to visualize the actin microfilaments (a-f) and actin polymerization sites (g-h) in fixed (a-f) and living cells (g, h), respectively. Scale, 20 µm.

### Pharmacological and genetic restoration of root hair morphology

Aldolase1 is the first enzyme in the biosynthetic pathway of chorismate, a precursor of AAA. Therefore, we evaluated if the altered root hair morphology and growth in the *ald1*-1 and *ald1*-2 plants were linked to a deficiency of AAA levels. The root hair development was analysed in 10 dpg seedlings of the different genotypes, germinated and grown on MS medium supplemented with an equimolar concentration of phenylalanine, tryptophan, and tyrosine. Root hairs with a restored tubular shape were observed in the two mutant alleles grown with a 50 and 100 µM mixture of AAA (Fig. 5a-c and Fig. S4). Interestingly, this effect was only present in the root tissue that was in contact with the agar that contained the cocktail of amino acids (Fig. 5a-c). To elucidate whether phenylalanine, tryptophan or tyrosine were separately contributing to prevent the altered root hair morphology and growth in the *adl1*-1 and *ald1*-2 mutants, the individual amino acids were added to the growth medium at 50 and 100 µM. The root hair swelling was prevented in both mutant alleles grown in medium supplemented with each amino acid, although to different extents. The percentage of plants with restored tubular root hair structure was 68–86 % with tyrosine, 36–77 % with phenylalanine, and 38–61% with tryptophan (Fig. S4). As described earlier, besides the swelling, the root hair growth is also arrested in the *aldolase1* mutants. In this regard, the maximal root hair length in the *adl1*-1 and *ald1*-2 mutants was not restored to the w. t. levels by adding different combinations and concentrations of AAA to the medium (Fig. 5d; Fig. S4). This result suggests that root hair structure and growth are mechanisms independently regulated by AAA. Interestingly, the root hair growth of Gifu plants grown in media supplemented with AAA was also affected (Fig. 5d), reinforcing the notion that root hair growth is influenced by AAA.

**Fig. S4.**
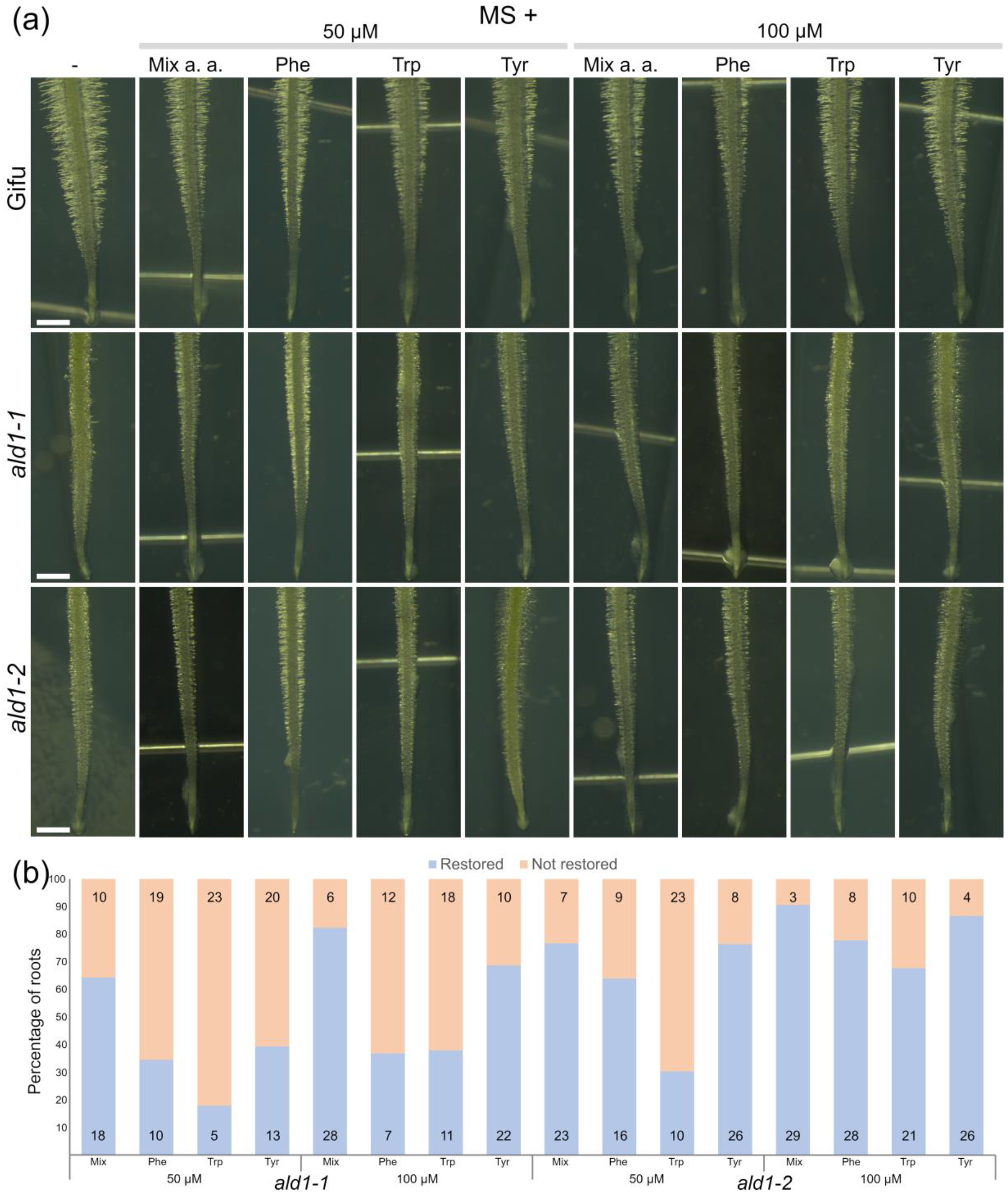
Phenotype of *Lotus* roots grown in medium supplemented with aromatic amino acids. (a) representative images of Gifu, *ald1*-1, and *ald1*-2 roots grown in MS medium supplemented individually with 50 or 100 µM of Phe, Trp, Tyr or a mixture of them. Scale, 1 mm (images captured at the same magnification). (b) proportion of plants with restored root hair morphology (tubular shape). The number of plants tested is indicated within the bars.

**Fig. 5.**
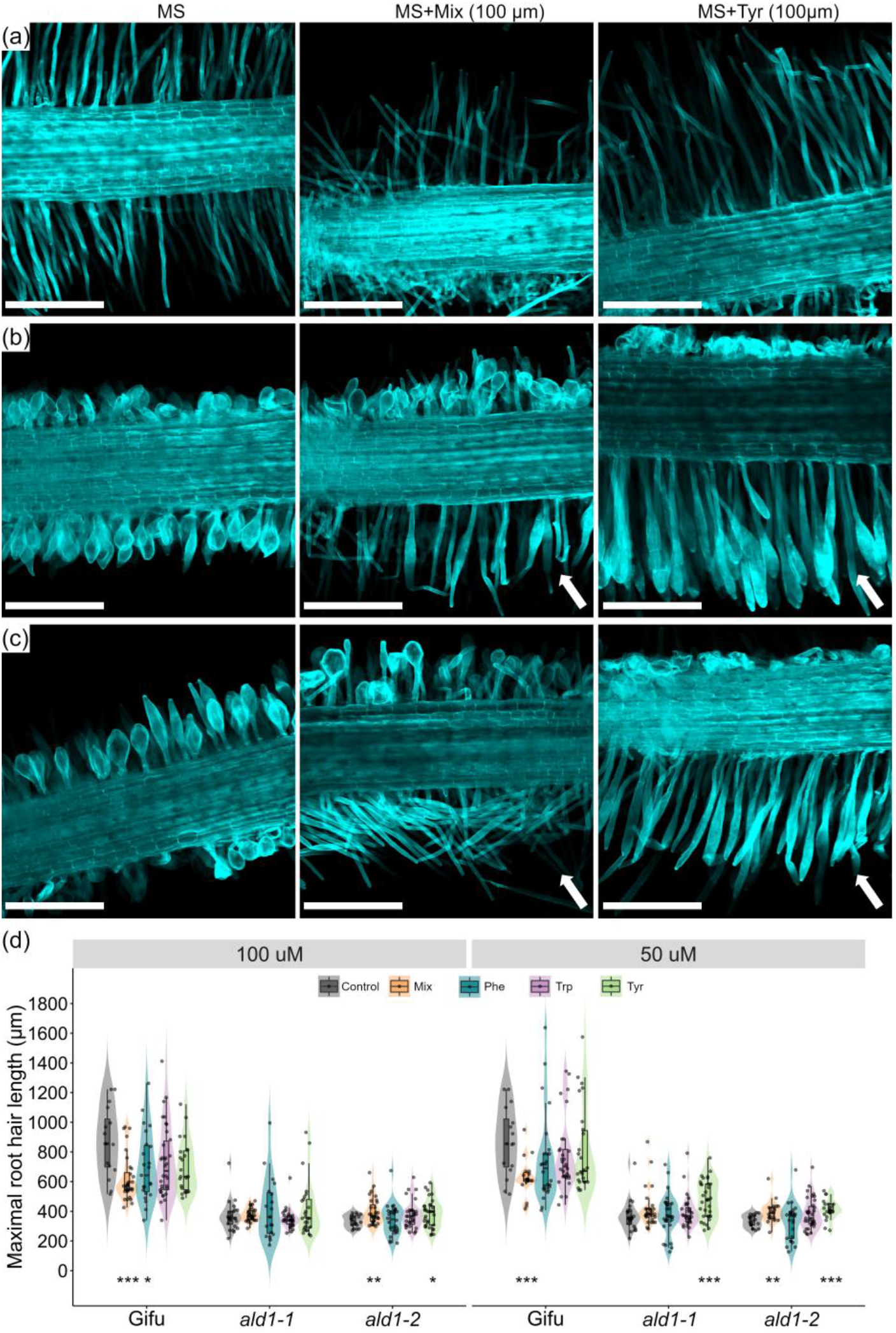
Restoration of root hair morphology in the *ald1*-1 and *ald1*-2 mutants. Confocal microscopy images of Gifu (a), *ald1*-1 (b) and *ald1*-2 (d) roots grown in MS medium (left panel) supplemented with Tyr (100 µM, right panel) or an equimolar combination of Phe, Trp and Tyr (100 µM, middle panel). The root hair morphology was restored only in the root tissue that was in contact with the agar (indicated with arrows). Scale, 500 µm. (d), violin dot plots show the maximal root hair length recorded in Gifu, *ald1*-1 and *ald1*-2 grown in MS medium supplemented with various concentrations of Phe, Trp and Tyr. Violin boxplots: centre line, median; box limits, upper and lower quartiles; whiskers, 1.5× interquartile range; points, individual data points. The number of plants tested are indicated in Fig. S4b. Student’s t-test of root hair length between Gifu and the two *aldolase1* mutant alleles. *p < 0.05; **p < 0.01; ***p < 0.001.

Using a chemical approach, we found that an imbalance in AAA levels was linked to the root hair phenotype in the *ald1*-1 and *ald1*-2 mutants. To demonstrate that this deficiency was caused by the disruption of the *LjAldolase1* gene, we made a construct composed of its coding sequence fused to the YFP transcriptionally regulated by the native promoter (*pAldolase1::Aldolase1-YFP*). The root hair morphology and development was restored in *Ljald1*-2 transgenic roots expressing the *pAldolase1::Aldolase1-YFP* construct (Fig. 6). Additionally, a clear fluorescent signal of the reporter was detected throughout the root hair development and other root tissues, supporting our previous findings. The fluorescence was associated to plastid-like structures, that correlates with the presence of a plastid signal peptide in the predicted amino acid sequence of *LjAldolase1*.

**Fig. 6.**
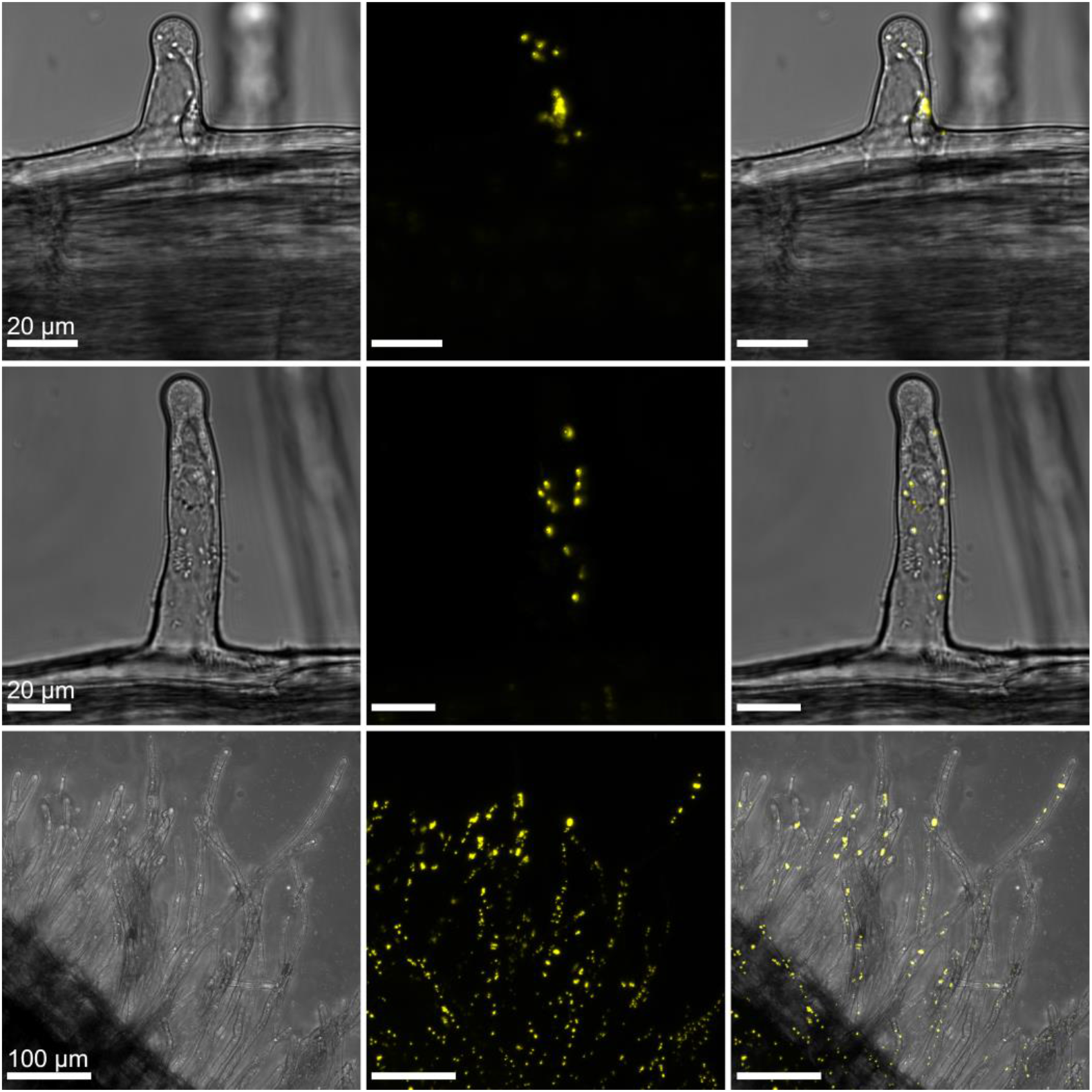
Subcellular localization of *LjAldolase1* and genetic complementation of *ald1*-2. Confocal microscopy images of *Ljald1*-1 transgenic roots transformed with the pAldolase1::Aldolase1-YFP construct. Left panel, transmitted light; Middle panel, yellow fluorescence; Right panel, merged images.

### Gene expression network associated with *LjAdolase1*

The evidence collected in this study demonstrates that disruption of *Aldolase1*, a gene involved in the biosynthetic pathway of AAA, has drastic consequences on root hair development. The data suggest that an insufficiency of these amino acids likely impacts the expression profile of genes and biological processes intrinsically connected to these molecules. We addressed this hypothesis by analysing the transcriptome of *ald1*-2 roots by RNAseq in 5 dpg seedlings. Compared with the root expression profile in Gifu plants of the same age, a total of 416 differentially expressed genes (DEG; p-adjust <0.5 and Log2FC ≥2) were detected (Fig. S5a). From this list, 163 sequences were upregulated and 253 repressed in *ald1*-2. Nineteen downregulated genes were related to the regulation of cell wall biomechanics. Interestingly, the symbiotic genes *CBS, Vapyrin1* and *Nin*, were significantly upregulated in the uninoculated *ald1*-2 roots, compared to Gifu (Fig. S5b).

**Fig. S5.**
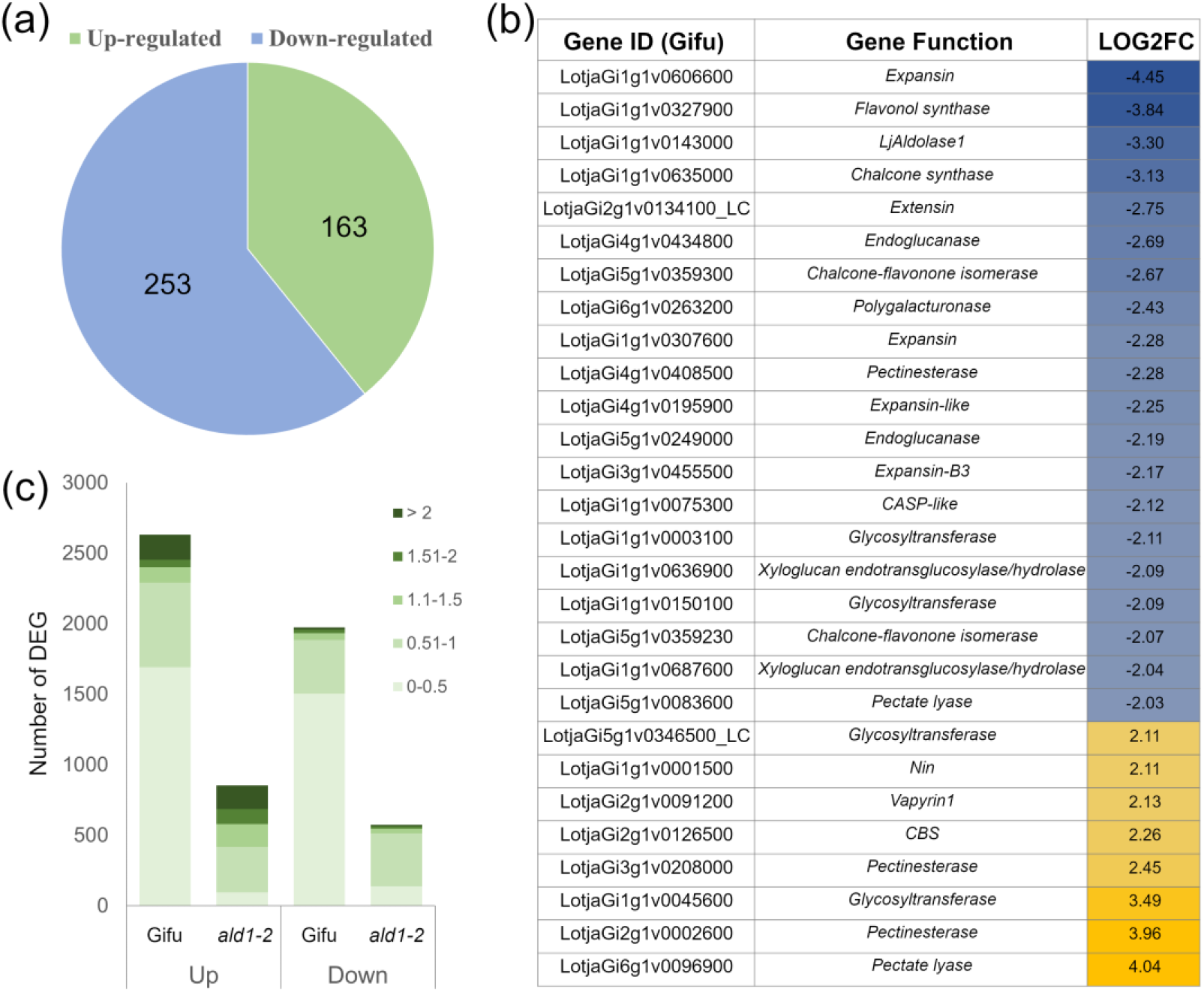
DEG in uninoculated 5 days old roots of *ald1*-2 mutant. (a), total number of up/downregulated genes (p-adjust < 0.5, Log2FC ≥2) respect to uninoculated roots of similar age in w. t. plants. (b), DEG encoding cell wall proteins, symbiotic genes and *LjAldolase1* in the RNAseq data of uninoculated roots in the *ald1*-2 mutant. (c), number of DEG in Gifu and ald1-2 roots at 5 dpi with IRBG74 (p-adjust < 0.5).

We previously described that *Aldolase1* was induced in Gifu roots at 5 dpi with IRBG74, when the major transcriptome response is observed in the intercellular infection (DEG Log2FC ≥2). To explore the contribution of *Aldolase1* to the symbiotic signalling in the *Lotus*-IRBG74 interaction, the expression pattern of *ald1*-2 roots was analysed at 5 dpi with IRBG74. Using low stringency parameters (p-adjust <0.5), we detected 2,840 DEG in Gifu roots and only 1,475 in *ald1*-2 (Fig. S5c). However, the number of DEG with Log2FC ≥2 was somewhat similar, 181 and 201 in *ald1*-2 and Gifu, respectively (Fig. 7a). Interestingly, less than 40% of these sequences overlapped (72), and none of the down-regulated genes overlapped (Fig. 7b,c). Despite the low similarity in the expression profiles between Gifu and *ald1*-2 roots (Fig. 7a-c), the main early symbiotic genes displayed comparable expression levels after IRBG74 inoculation (Fig. 7d).

**Fig. 7.**
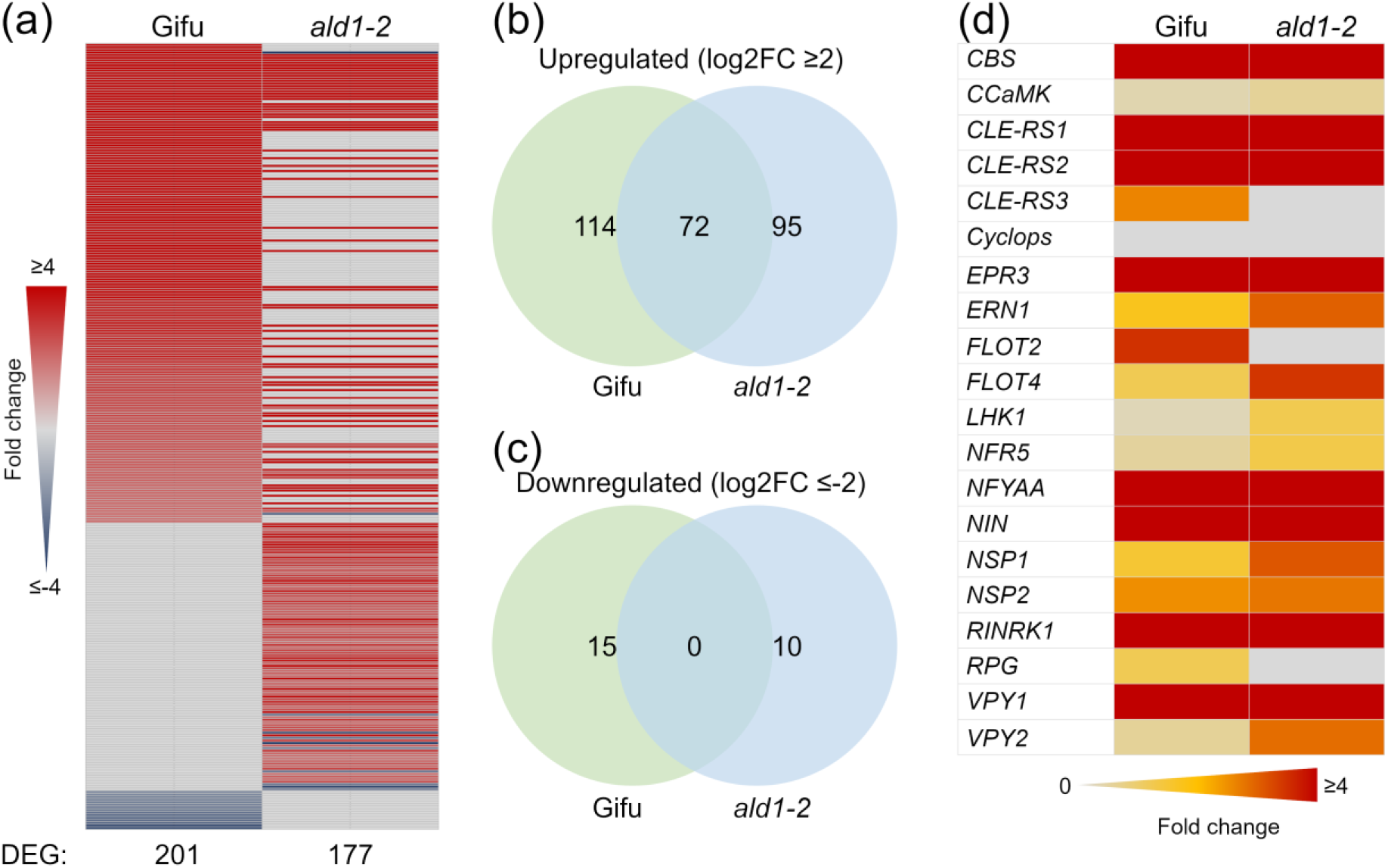
RNAseq expression of *ald1*-2 roots after IRBG74 inoculation. Heatmap (a) and Venn diagrams (b,c) of DEG in Gifu and *ald1*-2 roots at 5 dpi with IRBG74. (d), heatmap of early symbiotic genes in Gifu and *ald1*-2 (p-adjust < 0.5). The calculated values are respect to uninoculated roots of equivalent age on the same genotype.

### *LjAldolase1* contributes to rhizobial infection and nodule organogenesis

The RNAseq analysis in the *ald1*-2 roots inoculated with IRBG74 indicates that disruption of *LjAldolase1* affects the symbiotic transcriptome response. To evaluate the relevance of this finding, a nodulation kinetics analysis was conducted on plates for the *ald1*-1 and *ald1*-2 mutants, recording the pink and total number of nodules at 1 to 6 wpi with *M. loti* and IRBG74. Nodule formation (pink and total) was significantly reduced in both mutant alleles at most of the timepoints analysed, compared to Gifu (Fig. 8a-d; Fig. S6a,b). The delayed and reduced nodulation of *ald1*-1 and *ald1*-2 mutants had a negative influence on plant growth, since the length of the aerial parts was significantly shorter compared to Gifu at 6 wpi with any of the inoculum used (Fig. S6c,d). These findings reveal that *LjAldolase1* makes an important contribution to the nodulation capacity of *Lotus*. Since the root hair and nodulation phenotypes were consistent in the two mutant alleles, we decided to perform only a detailed symbiotic characterization for the *ald1*-2 allele.

**Fig. 8.**
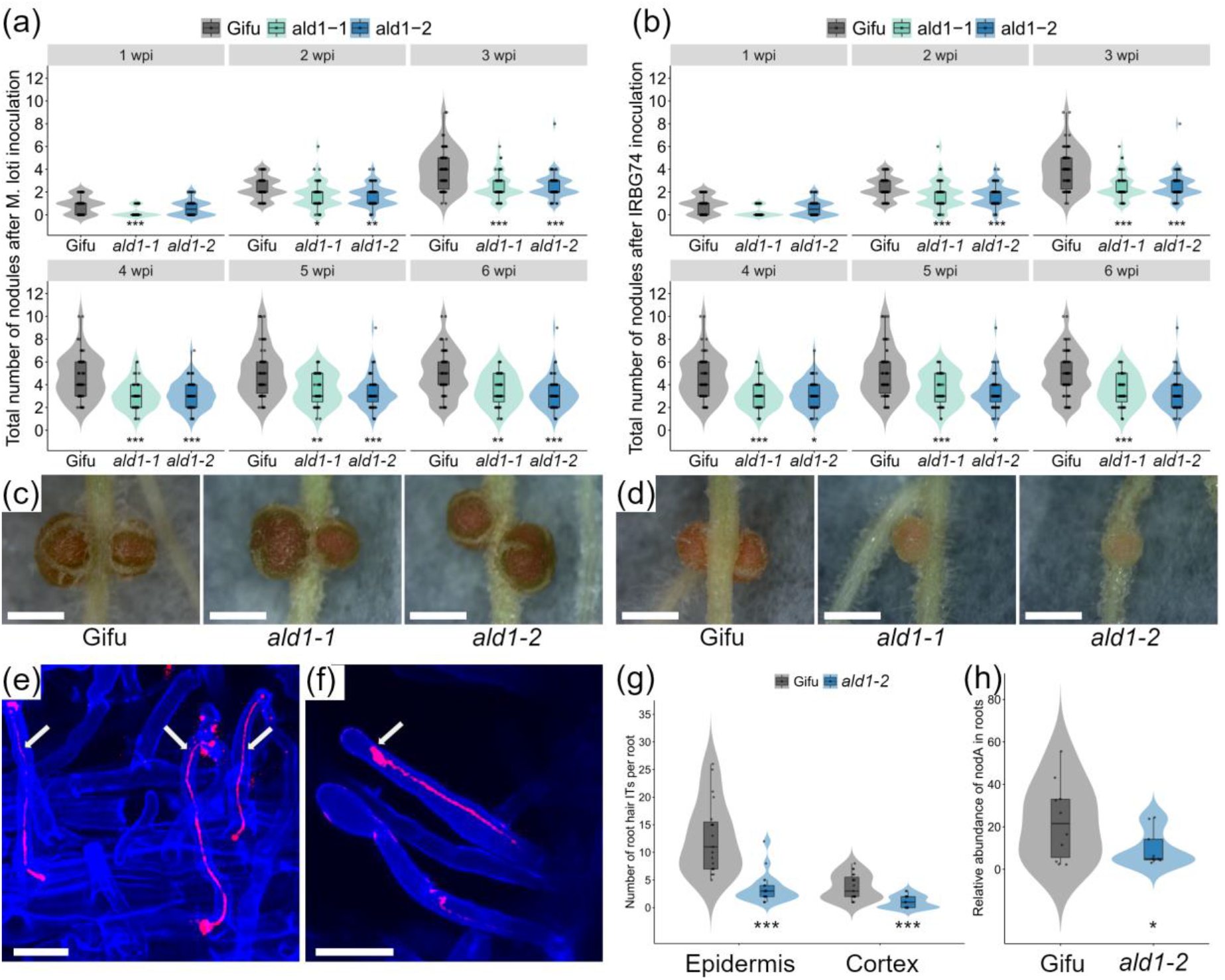
Intracellular and intercellular nodulation phenotype of *ald1*-1 and *ald1*-2 mutants. Total number of nodules recorded on Gifu (n ≥ 29), *ald1*-1 (n ≥ 43) and *ald1*-2 (n ≥ 29) at 1-6 wpi with *M. loti* (a) and IRBG74 (b). Mann–Whitney U-test of total number of nodules (asterisks below the violin graphs indicates significant difference: *p < 0.05; **p < 0.01; ***p < 0.001) between Gifu and the *aldolase1* mutant alleles. (c) and (d), representative images of nodules formed on different *Lotus* genotypes at 3 wpi with *M. loti* and IRBG74, respectively. Scale, 1 mm. Confocal microscopy images (e, f), and number of root hair ITs (g) at 1 wpi with *M. loti*-DsRed on Gifu (n= 19) and *ald1*-2 (n= 19). The arrows indicate the ITs. Scale bar, 20 µm. (h), abundance of IRBG74-nodA by qPCR in genomic DNA isolated from Gifu (n= 10) and *ald1*-2 (n= 9) roots at 3 wpi with IRBG74 and normalized to the LotjaGi1g1v0152000 gene accumulation. Student’s t test of *nodA* abundance between roots of Gifu and the mutants. *p <0.05 and ***p <0.001. Violin boxplots: centre line, median; box limits, upper and lower quartiles; whiskers, 1.5× interquartile range; points, individual data points.

First, the intracellular invasion of *M. loti*-DsRed was monitored on Gifu and *ald1*-2 roots by confocal microscopy. Root hair ITs were observed both in Gifu and *ald1*-2 at 1 wpi. In the latter, however, the typical root hair curling was not detected; instead, the IT initiation took place in a subapical region (Fig. 8e,f). Moreover, at this timepoint, the number of epidermal and cortical ITs *per* root was significantly lower in the *ald1*-2 mutant compared to Gifu (Fig. 8g). Since the intercellular infection of IRBG74 in *Lotus* is technically unsuitable for a quantitative evaluation through microscopy techniques, we followed a root endosphere approach; the *nodA* gene abundance was estimated by qPCR in DNA samples extracted from *Lotus* roots at 3 wpi with IRBG74 (Montiel et al. 2021). This analysis indicates that *ald1*-2 roots had a significant reduction in the relative abundance of the IRBG74-*nodA* gene, compared to Gifu (Fig. 8h).

**Fig. S6.**
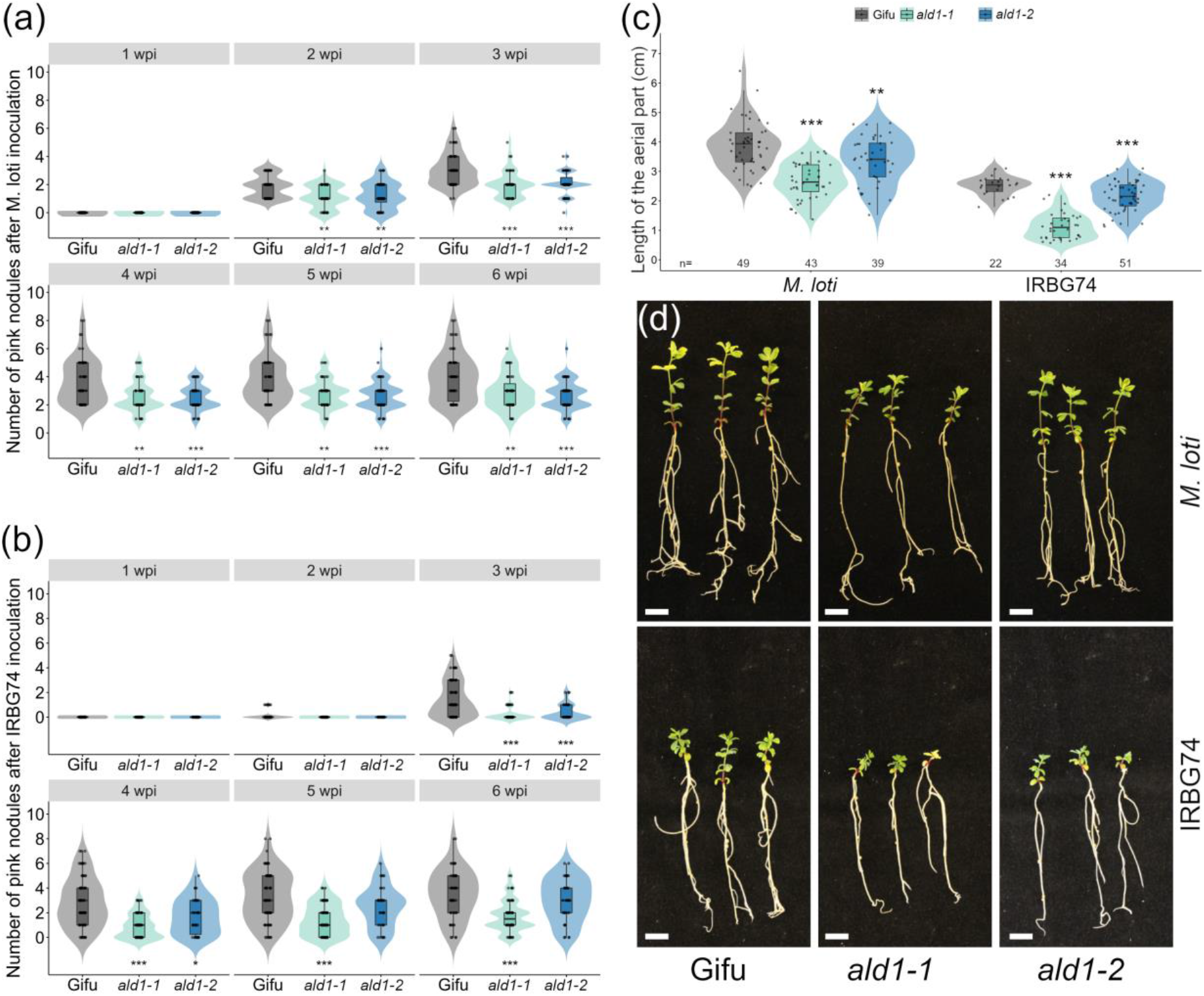
Nodule formation and plant growth in the *ald1*-1 and *ald1*-2 mutants. Violin dot plots show the number of pink nodules recorded on Gifu (n ≥ 29), *ald1*-1 (n ≥ 43) and *ald1*-2 (n ≥ 29) at 1-6 wpi with *M. loti* (a) and IRBG74 (b). Mann–Whitney U-test of pink nodules (asterisks below the violin graphs indicates significant difference: *p < 0.05; **p < 0.01; ***p < 0.001) between Gifu and mutant plants. Length of the aerial part (c) and representative images of Gifu, *ald1*-1 and *ald1*-2 plants (d) at 6 wpi with *M. loti* and IRBG74. Scale, 1 cm. Student’s t test of length of the aerial part between Gifu and the two *Aldolase1* mutant alleles inoculated with *M. loti* or IRBG74. **p < 0.01; ***p < 0.001. Violin boxplots: center line, median; box limits, upper and lower quartiles; whiskers, 1.5× interquartile range; points, individual data points.

Taken together, these results show that an efficient intra- and intercellular symbiotic infection in *Lotus* depends on the function of *Aldolase1*. These findings prompted us to investigate the bacteroid colonization in the nodules formed by *M. loti* and IRBG74 in the *ald1*-2 mutant. Infected nodule cells were detected in histological slides, stained with toluidine blue, of three-week-old nodules inoculated with *M. loti* and IRBG74 in Gifu (Fig. 9a,e) and *ald1*-2 (Fig. 9b,f). In Gifu nodules, the infected cells were densely packed with *M. loti* (Fig. 9a) and IRBG74 (Fig. 9e) bacteroids. By contrast, the infected cells in *ald1*-2 nodules were vacuolized with both inoculums, showing a deficient filling of the cells by bacteroids (Fig. 9b,f). Based on these findings, we decided to investigate the symbiosome structure by transmission electron microscopy (TEM). In Gifu nodules colonized by *M. loti*, the infected cells contained transcellular ITs with round-shaped symbiosomes hosting 1–2 bacteroids (Fig. 9c). Although the nodules in *ald1*-2 also contained 1 or 2 *M. loti* bacteroids per symbiosome, these organelle-like structures had irregular shapes. These abnormalities were also detected in the *ald1*-2 nodules formed by IRBG74, which were accompanied by an evident disruption in the space of the symbiosome (Fig. 9h), contrasting with the integrity of this structure observed in Gifu (Fig. 9g). Altogether these analyses thus showed that *Aldolase1* contributes to rhizobial infection and nodule organogenesis.

**Fig. 9.**
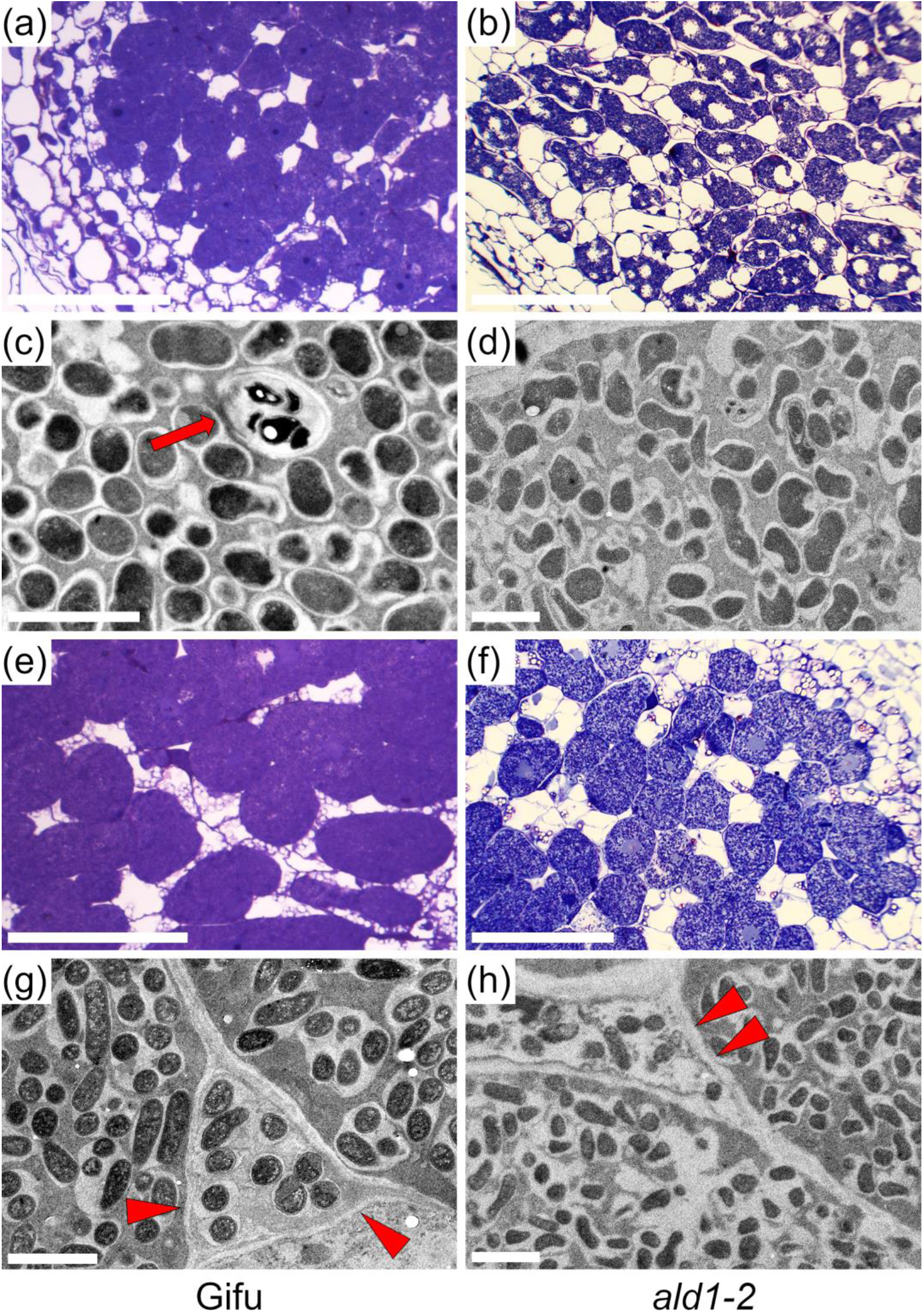
Nodule histology and symbiosomes in the *ald1*-2. Representative images of nodule sections and symbiosomes at 3 wpi with *M. loti* (a, c, e, g) and IRBG74 (b, d, f, h) obtained by transmitted light (a, b, e, f) and transmission electron microscopy (c, d, g, h) in Gifu (left panels) and *ald1*-2 (right panels). Arrow and arrowheads indicate transcellular infection threads and intercellular peg-infection, respectively. Scale, 2 µm (c, d, g, h) and 50 µm (a, c, e, g).

### AM colonization is delayed in *ald1*-2 roots

The expression profile of *LjAldolase1* during the RNS reflects its relevant role in this mutualistic association. Likewise, the data collected from the LjGEA show that *Aldolase1* is highly expressed in mycorrhized roots (Fig. S1b). This prompted us to evaluate the AM phenotype of *ald1*-2. For this purpose, 5 dpg seedlings of Gifu and *ald1*-2 were inoculated with *Rhizophagus intraradices* spores in Magenta boxes. Root fragments of Gifu and *ald1*-2 roots were collected at 4 wpi and stained with WGA-Alexa Fluor to visualize AM colonization by confocal microscopy. Fully branched hyphae with arbuscules within the cortex cells were observed in both genotypes, but the rate of arbuscule formation was apparently lower in the *ald1*-2 mutant compared to Gifu (Fig. 10a,b). Consequently, we conducted a quantitative analysis of different fungal colonization structures by optical microscopy in root segments stained with Trypan blue at 4 and 6 wpi (Trouvelot *et al*., 1986). Gifu and *ald1*-2 roots showed a similar percentage of mycorrhizal frequency at 4 and 6 wpi; a parameter that reflects the presence of AM fungi in the roots (Fig. 10c). However, other indicators that reveal the degree of fungal colonization were significantly reduced at 4 wpi in *ald1*-2: cortex mycorrhizal intensity, intensity of mycorrhiza in the root fragments, arbuscule abundance in mycorrhizal parts of root fragments and arbuscule abundance in the root system (Fig. 10c). These results show a delay in the establishment of AMS in *ald1*-2 plants, since the values obtained for all these parameters were similar to Gifu at 6 wpi (Fig. 10c). These results suggest that *Aldolase1* is required for the correct timing of AM root colonization.

**Fig. 10.**
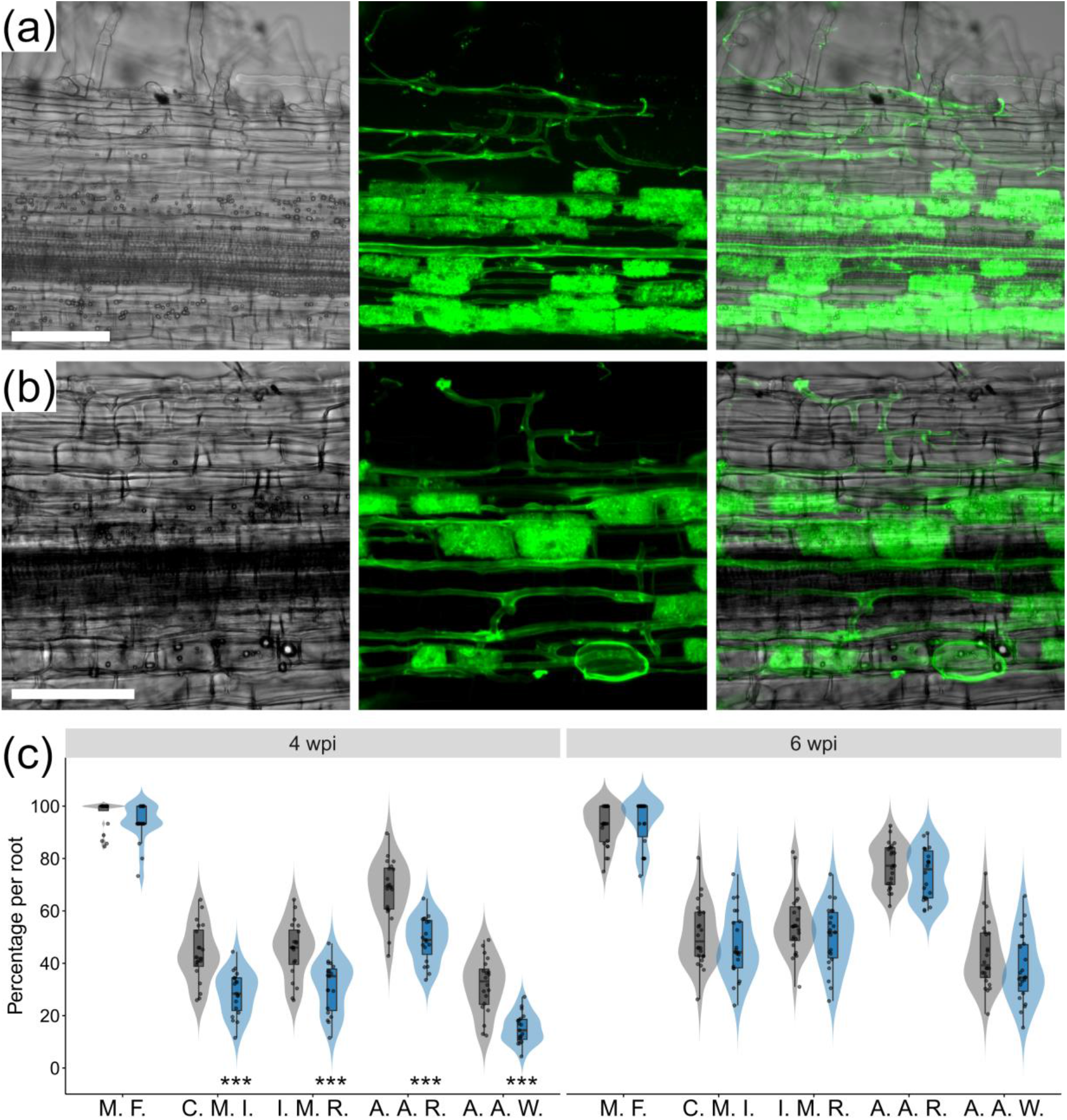
Arbuscular mycorrhization in the *ald1*-2 mutant. (a), mycorrhizal frequency (M. F.), cortex mycorrhizal intensity (C. M. I.), intensity of mycorrhiza in the root fragments (I. M. R.), arbuscule abundance in mycorrhizal parts of root fragments (A. A. R.) and arbuscule abundance in the root system (A. A. W.) of Gifu (n ≥ 20) and *ald1*-2 (n ≥ 19) plants at 4 and 6 wpi with *Rhizophagus irregularis*. Visualisation by confocal microscopy of AM colonisation at 4 wpi in Gifu (b) and *ald1*-2 (c) root fragments stained with WGA-Alexa Fluor 488. Left panel, transmitted light; Middle panel, green fluorescence; Right panel, merged images. Scale, 100 µm.

## Discussion

### Root hair development is AAA-dependent

Root hair biology is important and relevant due to the role played by these tubular extensions of epidermal cells on nutrient acquisition and its particularly polarized growth. The study of mutants defective in root hair emergence and growth shows that this process is highly complex and dynamic, involving transcription factors, phytohormones, cytoskeleton rearrangement, cell wall modifications, and secondary messengers (Shibata & Sugimoto, 2019). Our study demonstrates that perturbance of AAA biosynthesis in the *ald1*-1 and *ald1-*2 mutants influences the mechanism of root hair development. The observed progressive alteration in the root hair morphology of *ald1*-1 and *ald1*-2 mutants was prevented through genetic complementation with the *LjAdolase1* sequence, the first enzyme in the biosynthesis of AAA.

Besides the genetic complementation, we demonstrated that root hair swelling in *ald1-*1 and *ald1*-2 can be partially prevented in 80–90% of the plants by adding a mixture of AAA in a dose-dependent manner. This result indicates that in the *aldolase1* mutant alleles, the root hair phenotype was caused by the lack or deficiency of these aromatic molecules, and not by the absence of the intermediate compounds produced in the shikimate pathway. Interestingly, the tubular shape of root hairs was recovered in 68–86% of the mutants grown in a medium supplied only with 100 µM of tyrosine. This finding suggests that the root hair swelling in the *ald1*-1 and *ald1*-2 mutants was mostly provoked by the absence of tyrosine or its derived compounds. Phosphorylation of tyrosine residues occurs in actin-related proteins (Guillen *et al*., 1999), affecting the cytoskeleton dynamics in diverse biological processes such as plant bending and pollen tube growth (Kameyama *et al*., 2000; Zi *et al*., 2007). We observed that both root hair morphology and the actin cytoskeleton progressively deteriorated in *ald1*-1 and *ald1*-2. These effects were apparently preceded by changes in the distribution of F-actin plus ends. In healthy growing root hairs, the F-actin plus ends are localized at the root tip, paving the way for its growth (Zepeda *et al*., 2014). By contrast, these structures were detected in subapical regions of the root hairs in the *aldolase1* mutant alleles. The critical role played by the actin cytoskeleton during root hair development has been further supported by genetic evidence. The maintenance of the tip growth is prevented in the root hairs of the *Arabidopsis* mutant *deformed root hairs 1*, affected in the major actin of the vegetative tissue (Ringli *et al*., 2002). In *Lotus*, the actin cytoskeleton is severely compromised in the *actin-related protein component* (*arpc1*) mutant, which develops short and deformed root hairs (Hossain *et al*., 2012). Our data, along with previous reports, indicate that tyrosine has a strong influence on the cytoskeletal dynamics of plant cells and that the interference of its metabolism could impact the cell architecture.

The wide bulbous swelling of root hairs in the *ald1*-1 and *ald1*-2 mutants resemble the phenotype of the *root hair deficient 1* mutant in *A. thaliana*, disrupted in the UDP-D-glucose 4-epimerase (UGE) enzyme, necessary for the galactosylation of cell wall components (Seifert *et al*., 2002). Cell wall composition and flexibility is certainly crucial to sustain the root hair shape and growth, since the list of cell wall related mutants with defective root hairs includes the *leucine-rich repeat extensins Atlrx1* and *ATlrx2* (Baumberger *et al*., 2003), and the *cellulose synthases-like D* (Wang *et al*., 2001; Karas *et al*., 2021). L-phenylalanine and L-tyrosine are products of the shikimate pathway and also serve as precursors for the phenylpropanoid metabolism. These secondary metabolites include flavonoids and cell-wall associated phenolics (Vogt, 2010). It was recently shown that flavonols modulate the reactive oxygen species (ROS) levels that drive root hair development in *A. thaliana*. Mutants affected in synthesis of flavonols exhibit a greater frequency of trichoblast cells forming root hairs and raised epidermal ROS levels (Gayomba & Muday, 2020). Our RNAseq analysis of *ald1*-2 roots suggests that the root hair phenotype in *ald1*-2 could be related to a miss regulation of several genes related to cell wall biomechanics such as: *Expansin, Pectate lyase, Xyloglucan endoglucanase, CASP-like protein, Endoglucanase* and *Pectinesterase*.

### *Aldolase1*, a player in the common symbiotic response

In the *Lotus*-rhizobia symbiosis, *Aldolase1* expression is clearly linked to nodule formation, since its promoter was strongly expressed in developing nodules and the number of these structures was significantly reduced in the *ald1*-1 and *ald*-2 mutants after *M. loti* or IRBG74 inoculation. This symbiotic phenotype is probably caused by insufficient levels of AAA and their derived compounds, in a high-demanding organogenesis program (Mergaert *et al*., 2020). *Aldolase1* is apparently an integral component of the genetic program governing developmental processes, since its promoter was also highly active in the root apical meristem and emerging lateral roots. In *M. truncatula*, it was recently shown that a large proportion of the transcriptome changes in lateral root primordia also occur in developing nodules (Schiessl *et al*., 2019). Additionally, the delay in the nodulation kinetics observed in the *aldolase1* mutant alleles could be provoked by deficient reprograming of the transcriptome. The RNAseq analysis of *ald1*-2 roots inoculated with IRBG74 showed that, although the major symbiotic genes were induced, the total number of DEG was considerably lower compared to the transcriptome response in Gifu wild type and only a minor proportion of DEG overlapped between these two genotypes. *Aldolase1* is the major Phospho-2-dehydro-3-deoxyheptonate aldolase isoform expressed in *Lotus* roots; therefore, it is likely that disruption of this gene has a negative impact on the flavonoid levels derived from phenylalanine. Flavonoids are essential compounds produced by legume roots in the rhizosphere for their chemical crosstalk with rhizobia (Liu & Murray, 2016), and silencing of flavonoid biosynthesis components inhibits nodulation in *Glycine max* and *M. truncatula* transgenic roots (Subramanian *et al*., 2006; Wasson *et al*., 2006). Disruption of *LjAldolase1* also interferes with rhizobial colonization both intra- and intercellularly. The presence of IRBG74 was reduced by 50% in *ald1*-2 roots compared to Gifu, at 3 wpi. However, the impact on the intracellular infection by *M. loti* was more dramatic, as the number of epidermal and cortical root hair ITs diminished by >70% in *ald1*-2 at 1 wpi compared to Gifu. This pronounced effect is likely influenced by the abnormal and collapsed root hairs in the *aldolase1* mutants, being unable to host an IT. Additionally, root hair curling, a key structure to trap rhizobia and initiate IT formation, was not observed in *ald1*-2 plants. Interestingly, the *Aldolase1* promoter was not detected in the root hairs infected by *M. loti* or in the twisted and curled root hairs induced by IRBG74, but rather in the root cells surrounding the infection sites.

The analysis of legume mutants indicates that the genetic requirements for RNS and AMS are partially overlapping, leading to the concept of a common symbiosis signalling pathway (CSSP) (Parniske, 2008). Recent phylogenomic studies confirm that certain common symbiosis genes have been retained in different plant lineages that engage intracellular symbiotic associations (Radhakrishnan *et al*., 2020). However, several transcriptome analyses in legumes show that a larger set of genes are induced in mycorrhized and nodulated roots (Manthey *et al*., 2004; Deguchi *et al*., 2007; Handa *et al*., 2015; Nanjareddy *et al*., 2017). For instance, mycorrhization and nodulation is affected in *Lotus* roots silenced or disrupted in the *Lectin Nucleotide Phosphohydrolases* (*LNP*), and the *SNARE* genes *LjVAMP72a* and *LjVAMP72b* (Roberts *et al*., 2013; Sogawa *et al*., 2018). We found that although the different fungal structures were formed in the *ald1*-2 roots at 4 wpi, their numbers were significantly lower than in Gifu. However, the proportion of the AM components in the root system was comparable to the w. t. at 6 wpi, reflecting a delay in the colonization process and suggesting a role of *Aldolase1* in timing of this process. Similarly, a milder symbiotic phenotype with AM fungi compared to the RNS has been also described for *ccamk* and *symrk* mutants in *Lotus* (Demchenko *et al*., 2004).

## Conclusion and perspectives

AAA are the building blocks of indispensable metabolites and proteins for cell functioning. Interestingly, no pleiotropic effects were observed in the *ald1*-1 and *ald1*-2 mutants, but instead specific phenotypes in root hair development and mutualistic associations. The other two *Aldolases* expressed in *Lotus* roots could supply the minimal requirements of the cell but were still insufficient for the aforementioned processes. The link of some AAA and derived compounds with the cell wall and actin cytoskeleton is consistent with developmental and symbiotic defects observed in the *aldolase1* mutant alleles. However, further research is needed to fully understand the specific metabolites and proteins involved in these different processes.

## Data availability

The RNAseq reads associated with this study are available in the SRA under bioproject accession number PRJNA632725.

## Acknowledgements

This work was supported by the grant Engineering the Nitrogen Symbiosis for Africa made to the University of Cambridge by the Bill & Melinda Gates Foundation (ENSA; OPP11772165), the European Research Council (ERC) under the European Union’s Horizon 2020 research and innovation programme (grant agreement no. 834221). Mexican Consejo Nacional de Ciencia y Tecnología (CONACyT, grant A1-S-9236) and Dirección General de Asuntos del Personal Acade′mico (DGAPA)-Universidad Nacional Autónoma de México (UNAM) – Programa de Apoyo a Proyectos de Investigación e Innovación Tecnológica (PAPIIT, grant IN204221).

## Author contributions

J.M. and J.S. conceived the research. J.M., E.K.J., I.G-S., S.N-M. and L.C. performed the experiments. D.R. processed the RNAseq data. J.G.D. and J.S. revised the manuscript. J.M. wrote the manuscript.

